# Stabilisation of β-Catenin-WNT signalling by USP10 in APC-*truncated* colorectal cancer drives cancer stemness and enables super-competitor signalling

**DOI:** 10.1101/2023.02.10.527983

**Authors:** Michaela Reissland, Oliver Hartmann, Saskia Tauch, Cristian Prieto-Garcia, Clemens Schulte, Daniel Solvie, Sinah Loebbert, Anne-Claire Jacomin, Marina Pesic, Jeroen M. Bugter, Christina Schuelein-Voelk, Carmina T. Fuss, Nikolet Pahor, Carsten Ade, Viktoria Buck, Michael Potente, Vivian Li, Gerti Beliu, Armin Wiegering, Eliya Bitman-Lotan, Tom Grossmann, Mathias Rosenfeldt, Martin Eilers, Hans Maric, Madelon M. Maurice, Florian Greten, Ivan Dikič, Amir Orian, Peter Gallant, Markus E. Diefenbacher

## Abstract

The contribution of deubiquitylating enzymes to β-Catenin stabilisation in intestinal stem cells and colorectal cancer (CRC) is poorly understood. Here, we report the deubiquitylase USP10 as an APC-truncation- specific enhancer of β-Catenin stability, potentiating WNT signalling in CRC and cancer stem cells. Mechanistically, interaction studies in various CRC cell lines and in vitro binding studies, together with computational modelling, revealed that USP10 binding to β-Catenin is mediated via the unstructured N-terminus of USP10 and requires the absence of full-length APC. Notably, loss of USP10 in CRISPR engineered intestinal organoids reduces tumorigenic properties of CRC and blocks the super competitor-signalling of APC-mutated CRC. Furthermore, reduction of USP10 induces the expression of differentiation genes, and opposes the APC-truncated phenotype in an intestinal hyperplasia model of *D.melanogaster*.

Taken together, our findings reveal USP10s role in intestinal tumourigenesis by stabilising β-Catenin, leading to aberrant WNT signalling, enhancing cancer cell stemness and implicate the DUB USP10 as a cancer specific therapeutic vulnerability in *Apc* truncated CRC.

## Introduction

Colorectal cancer (CRC) is the third most common cancer for both sexes, with 8 % estimated new cases and 8 to 9 % estimated deaths in 2021 (Sung et al., 2021). It is widely accepted that environmental risk factors, such as diet, obesity, low level of exercise, alcohol and tobacco consumption increase the risk of developing CRC. Besides these environmental risk factors, inherited forms of CRC, characterized by distinct mutations, occur at a lower frequency (Half et al., 2009). Although most patients have an overall good survival after diagnosis, only 10 % patients with progressed disease survive longer than 5 years (Siegel et al., 2021). This poor prognosis for late-stage patients punctuates the necessity of revealing novel and exploitable vulnerabilities in CRC.

In 80 % of cases, CRC is characterized by hyperactivation of WNT signalling (Novellasdemunt et al., 2015). This is predominantly caused by truncating mutations in the tumour suppressor gene *Adenomatous Polyposis Coli* (*APC*). These truncating mutations are causative to the impaired ability of the WNT destruction complex to degrade the WNT effector β-Catenin (Novellasdemunt et al., 2015). Several ‘hotspot’ mutations have been described for *APC*, including the catenin inhibitory domain (CID), resulting in various variants of APC truncation identified in patients (Ranes et al., 2021). This domain contains the β-Catenin binding 20 amino acid repeats (20 AAR). Consequently, *APC*-truncated tumour cells accumulate the co-activator β-Catenin, which translocates to the nucleus, displaces the TLE co-repressor and associates with the transcription factor TCF-4/LEF-1 to promote the expression of WNT pathway target genes (Morin et al., 1997; Yang et al., 2006). The accumulation of β-Catenin results in elevated cellular proliferation independent of WNT ligand availability and is believed to be a major event in disease onset of CRC (Zhan et al., 2017; Zhao et al., 2022).

Truncating mutations within *APC* impact β-Catenin abundance or activity and subsequently oncogenic transformation (Barua and Hlavacek, 2013; Hankey et al., 2018; Novellasdemunt et al., 2017; Yang et al., 2006) (Schneikert et al., 2007), as these cells only exhibit a limited ability to ubiquitylate β-Catenin (Ranes et al., 2021). Despite a basal post transcriptional modification (PTM) activity retained by truncated APC (Ranes et al., 2021), several E3 ligases were described in the past to modulate β-Catenin ubiquitylation independent of the WNT-destruction complex (Chen et al., 2018; Gao et al., 2014), including, but not limited to, HUWE1 (Dominguez-Brauer et al., 2017), FBXW7 (Jiang et al., 2016), JADE-1 (Chitalia et al., 2008), UBR5 (Shearer et al., 2015) and RNF4 (Thomas et al., 2016).

Apart from ubiquitin ligases, several deubiquitylating enzymes (DUBs) were reported to deubiquitinate β-Catenin (Park et al., 2020), such as USP7 (Novellasdemunt et al., 2017) or USP20 (Wu et al., 2018), thereby extending the regulation of β-Catenin activity or abundance. However, whether the APC truncation status impacts the stabilization of β-Catenin by DUB enzymes is unkown. The possibility arises that truncated APC not only prevent the physiological degradation of β-Catenin, but also serves as a scaffold for β-Catenin regulating DUBs, enhancing β-Catenin accumulation in CRC, in an APC truncation specific manner. We investigated whether partial loss of discreet domains within APC allow *de novo* protein-protein interactions, towards which tumours could become addicted.

Here, we identified the DUB Ubiquitin Specific Protease 10 (USP10) as a direct binder of β-Catenin. The protein interaction between the DUB and the WNT effector only occurs in cells which have lost all AAR domains within *APC*, mutations present in around 30% of all CRC patients (Cerami et al., 2012; Gao et al., 2013). USP10 is crucial in the tumorigenic signalling of β-Catenin and its abundance strongly enhanced WNT target gene expression and is required to maintain a *stemness-like* expression profile, in CRC and murine tumour organoids, respectively. Moreover, this mechanisms is evolutionarily conserved as elimination of Usp10, suppressed the gut progenitor hyperproliferation phenotype and reduced survival in homozygous *Apc* (*Apc*^*Q8/Q8*^) truncation in a *D*.*melanogaster* model (Ahmed et al., 1998).

Overall, our study demonstrates that USP10 stabilises β-Catenin, promotes WNT-signalling and cancer stemness, essential to tumour onset and development in an *APC*Δ^*AAR*^-truncation dependent manner. Taken together our results suggest that targeting USP10 could thus be a viable treatment option for the large cohort of CRC patients carrying *APC*Δ^*AAR*^-truncating mutations.

## Results

### Identification of USP10 as a novel regulator of β-Catenin signalling in CRC

In colorectal cancer, increased abundance of the WNT effector β-Catenin is mediated either by loss of function mutations and truncations within the *APC* gene or via mutations of the degron motive within *CTNNB1*, the gene encoding β-Catenin. To investigate if these alterations impact the ubiquitylation of β-Catenin in CRC, we conducted an endogenous ubiquitin TUBE (tandem ubiquitin binding entity) assay in a panel of human CRC lines, comprising β-Catenin mutant lines (HCT116 and LS174T), or cell lines varying in the truncation length of APC, DLD-1, SW480, SW620, Colo320 and HT-29, respectively (Figure 1A). Remarkably, and irrespective of genetic alteration we were able to detect poly-ubiquitylation of β-Catenin (Figure 1A). This suggested that protein stability of the WNT effector could be altered in a Ubiquitin Proteasome System (UPS) specific fashion. To this end, we performed a human DUB siRNA screen in the APC truncated CRC line HT-29, followed by assessing the residual protein abundance of β-Catenin (Figure 1B, C). Analysis of the screen identified the deubiquitylase USP10 as a positive regulator of β-Catenin stability, along with previously identified DUBs (Figure 1C and S1A, B). Indeed, we verified that loss of USP10 resulted in the depletion of the cytosolic and nuclear pool of β-Catenin, as seen by immunofluorescence (Figure 1C).

**Figure 1:**
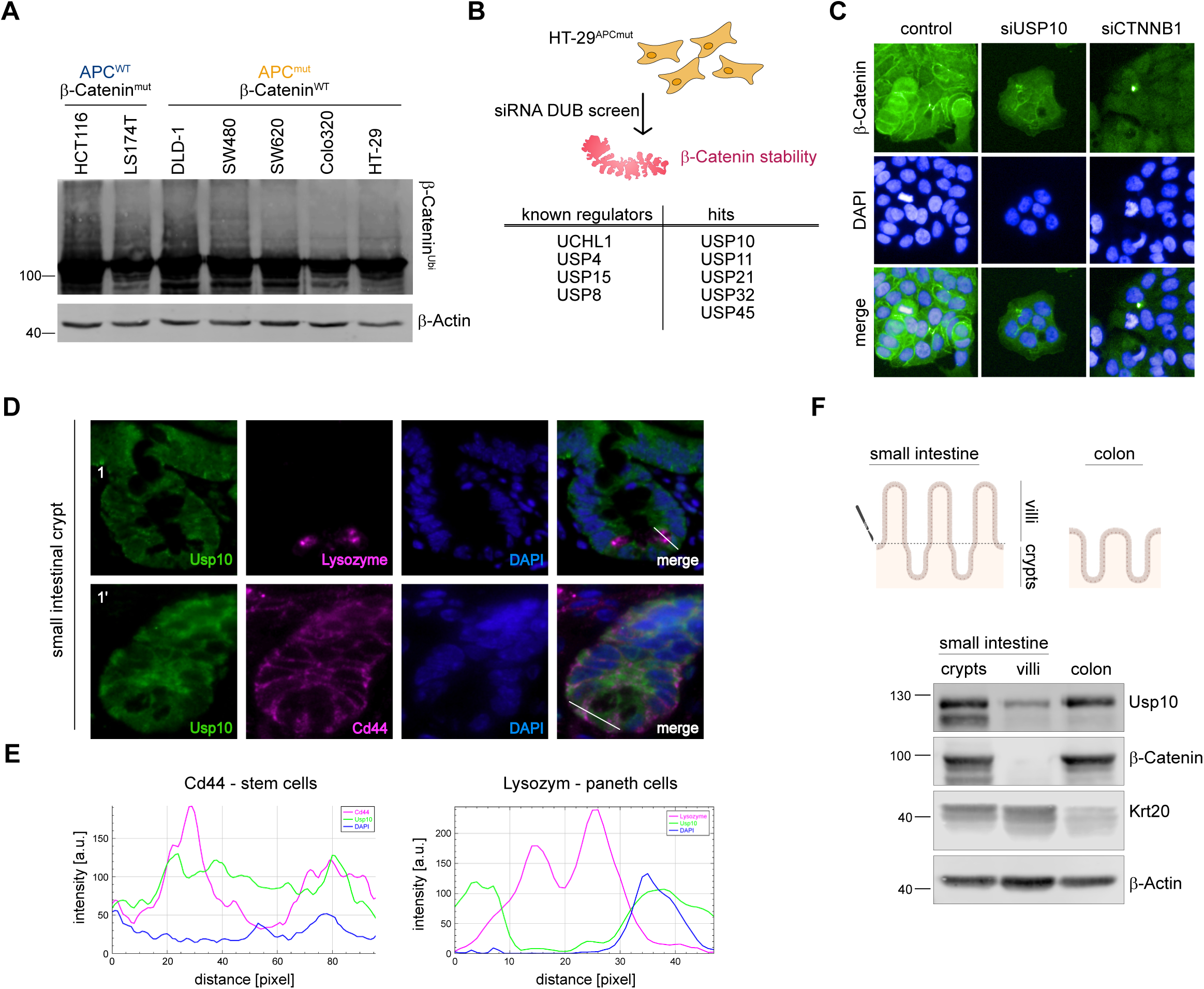
Identification of USP10 as a novel regulator of β-Catenin signalling in CRC. A. Tandem Ubiquitin Binding Entity (TUBE) assay of endogenous poly-ubiquitylated proteins, followed by immunoblotting against endogenous β-Catenin in human CRC cell lines with varying mutations. HCT-116 and LS174T mutant for β-Catenin, DLD-1, SW480, SW620, Colo320 and HT-29 mutant for *APC*. β-Actin served as loading control. B. Schematic model of siRNA DUB library screen conducted in *APC* mutant HT-29 cells. Cells were transfected with 4 individual siRNA against DUBs and 48 hours post transfection immunofluorescence against endogenous β-Catenin was analysed via Operetta high-content microscope imaging. n=3. DAPI served as nuclear marker. Identified known and new putative regulators of β-Catenin are highlighted. C. Representative immunofluorescent images of endogenous β-Catenin (green) upon siRNA mediated knock-down of NTC (control), *CTNNB1* and *USP10*, respectively. DAPI served as nuclear marker (blue). D. Representative immunofluorescent images of WT intestinal crypts of endogenous Usp10 (green) and crypt cell specific markers. Upper panel: Lysozyme (magenta) marks Paneth cells. Lower panel: Cd44 (magenta) labels stem cells. DAPI served as nuclear marker (blue). E. Histogram of fluorescence over indicated length in D. F. Schematic representation of murine small intestine and colon. Villi were scratched from the intestine and small intestinal and colonic crypts were isolated using EDTA. Isolated tissue from two individual mice was analysed for endogenous abundance of Usp10, β-Catenin and Krt20. β-Actin served as loading control. (n=2)

Since USP10 was not implicated in intestinal homeostasis and β-Catenin signalling, we investigated its abundance in murine intestine next (Figure 1D to F). Usp10 was highly abundant in intestinal crypts when compared to the villus (Figure S1C, D), and nuclear localised in intestinal stem cells (Usp10^+^/Cd44^high^; Usp10^+^/Lysozyme^-^; Figure 1D, E). A crypt specific expression of Usp10 was further validated using murine intestine fractionation immunoblotting of crypt versus villus samples, derived from small intestine or colon, respectively (Figure 1F).

These data propose a novel regulator of β-Catenin stability, USP10, and a putative involvement of this DUB in WNT signalling, intestinal homeostasis and carcinogenesis.

### USP10 is upregulated in CRC and correlates with poor patient survival

To further investigate whether the abundance of USP10 is enriched upon transformation of intestinal cells first, we performed histopathologic analysis of individual tumours in animals, where carcinogenesis was induced either by acute CRISPR editing of *Apc*, generating an endogenous truncated APC (APC^ex10^) or by loss of heterozygosity of *Apc* in *Apc*^*min/+*^ mice (Figure 2A-D, S2A-D). Irrespective of genetic alteration of *Apc* causal to carcinogenesis did we observe that the protein level of Usp10 was significantly upregulated in tumours within the GI tract, along with elevated protein levels of β-Catenin, when compared to non-transformed or non-oncogenic intestinal epithelium, respectively (Figure 2C, D and S2C-D).

**Figure 2:**
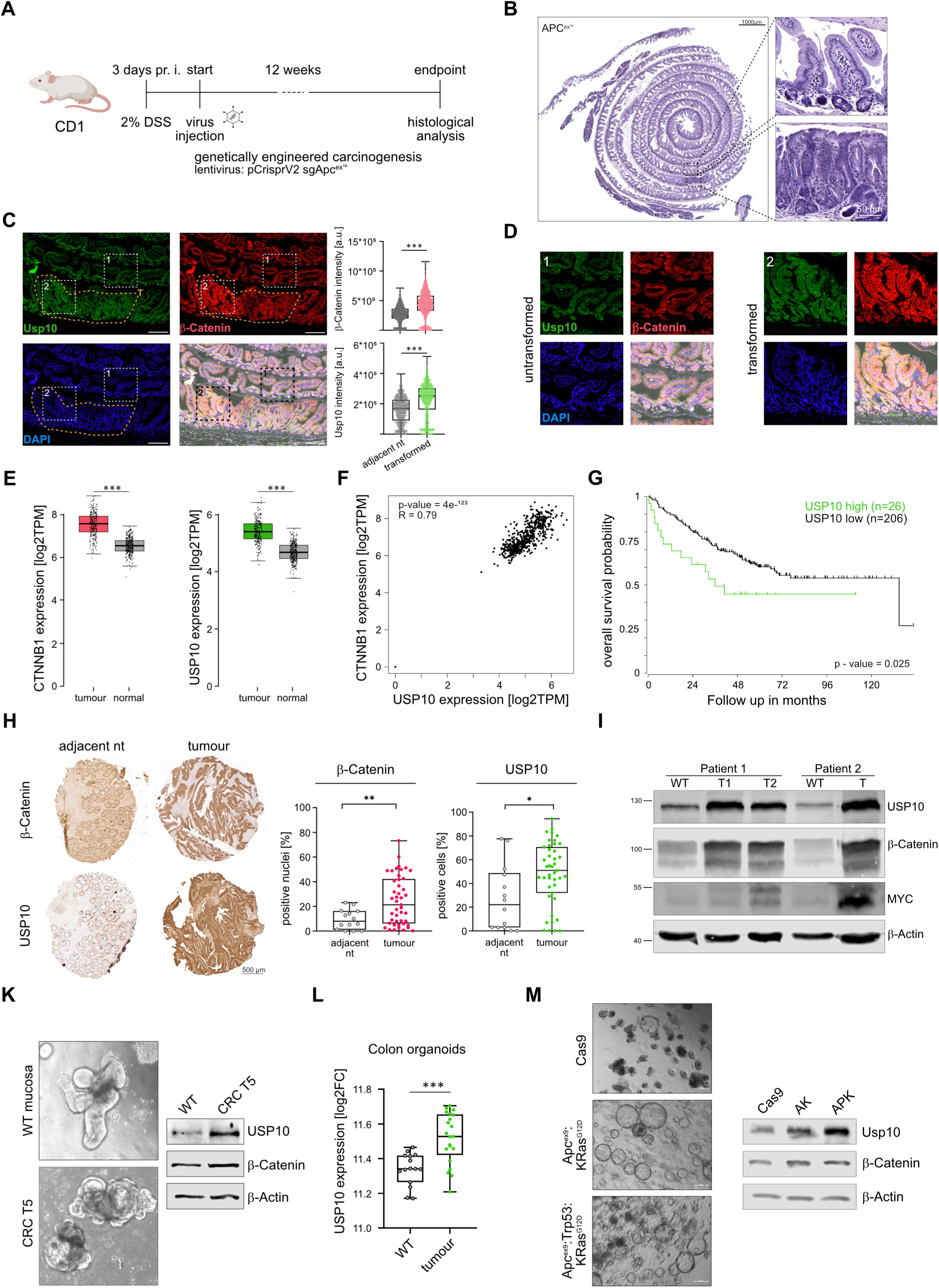
USP10 is upregulated upon oncogenic transformation. A. Schematic representation of acute *in vivo* CRC onset in wild type CD1 animals using colorectal instillation of lentivirus particles encoding sgRNA against murine *Apc*, targeting exon 10 (*Apc*^*ex10*^), and constitutive expression of SpCas9. Viral backbone was pLenti-CRISPR-V2. pr.i. -pre infection B. Haematoxylin and eosin (H&E) staining of CRISPR mediated tumour onset in CD1 animals, 12 weeks post intracolonic instillation of virus. Insets highlight either non-transformed adjacent tissue or primary tumour upon *Apc* deletion. C. Representative immunofluorescent images of mice shown in Figure 1F and G of endogenous Usp10 (green) and β-Catenin (red). DAPI served as nuclear marker (blue). Insets highlight either untransformed (1) or transformed (2) regions. Intensity of β-Catenin and Usp10 staining was quantified using QuPath software. P-values were calculated using Mann-Whitney test. Individual cells/values are highlighted as dots. ***p<0.001 D. Insets from Figure 1H. High magnification immunofluorescent images of intestines of CRISPR infected animals for Usp10 (green) and β-Catenin (red). DAPI served as nuclear marker (blue). E. Expression of *CTNNB1* and *USP10* in non-transformed (normal) and CRC (tumour) samples. Publicly available data from GEPIA. COAD (n=275) and GTEx (n=349) data were displayed as boxplots for *USP10* and *CTNNB1* expression. P-values were calculated using one-way ANOVA. Data was visualised using the online tool www.gepia.cancer-pku.cn. ***p<0.001. F. Correlation of gene expression between *CTNNB1* and *USP10* in human CRC. R: Spearman’s correlation coefficient. n^T^=275 n^N^=349. Data was visualised using the online tool www.gepia.cancer-pku.cn. G. Publicly available patient survival data of CRC patients are stratified by relative expression of *USP10*. n□=□206 (low) and n□=□26 (high). Survival correlation analysis was performed using R2: Genomics Analysis and Visualization Platform, using the Tumour Colon - Smith dataset. H. Representative images of immunohistochemistry (IHC) staining of a Tissue Micro Array (TMA) from CRC patients, comprising adjacent non-transformed tissue (adjacent nt) and CRC samples against β-Catenin and USP10. P-values were calculated using Mann-Whitney U test. *p<0.05; **p<0.005. I. Immunoblotting of endogenous abundance of USP10, β-Catenin and MYC in non-transformed (WT) and patient matched CRC tumour samples (T) from two individual patients. β-Actin served as loading control. K. Representative brightfield images of patient derived intestinal organoids, comprising either wild type (WT mucosa) or tumour derived organoids (CRC T5), respectively. Immunoblotting of endogenous abundance of USP10 and β-Catenin of patient organoids. β-Actin served as loading control. L. Expression of USP10 in non-transformed (WT) and CRC patient derived organoids (tumour). Analysis was performed using R2: Genomics Analysis and Visualization Platform, using the Organoid - Clevers dataset. P-values were calculated using Mann-Whitney U test. ***p <0.001. M. Representative brightfield images of murine intestinal organoids, comprising either wild type (Cas9), after targeting and growth factor depleted selection upon CRISPR engineering of *Apc* exon 9 (*Apc*^*ex9*^) and *KRas* to *KRas*^*G12D*^ *(A*^*ex9*^*K*^*G12D*^), or upon co-deletion of *Trp53 (A*^*ex9*^*PK*^*G12D*^), respectively. Immunoblotting of endogenous abundance of Usp10 and β-Catenin in Cas9, *AK* and *APK* organoids. β-Actin served as loading control.

Next, we determined expression level of USP10 by interrogating publicly available patient data of colorectal cancer (Cerami et al., 2012). While *USP10* was rarely mutated in CRC, it was predominantly upregulated, along with *CTNNB1*, when compared to adjacent wild-type tissue (Figure 2E and S2E), and *USP10* and *CTNNB1* demonstrated a significant degree of correlation of expression in CRC samples (Figure 2F, n^T^=275, n^WT^ = 349, Spearman coefficient R = 0.79). Remarkably, elevated expression of USP10 in CRC is a strong indicator of overall poor patient survival in CRC (Figure 2G). Prompted by this observation, we next studied USP10 and β-Catenin levels by Immunohistochemistry (IHC) of tissue micro arrays (TMA) of CRC patients comprising non-transformed and tumour tissue (Figure 2H). Not only was a difference in tissue architecture observed in CRC tissue samples, but USP10 and β-Catenin indeed showed a significant upregulation in CRC when compared to the adjacent tissue (Figure 2H). Elevated protein abundance was furthermore analysed by using human samples from curative CRC resection surgeries, subjected to immunoblotting against endogenous USP10, β-Catenin and MYC. Non-transformed, adjacent tissue served as control. USP10, β-Catenin and the oncogene MYC were increased in tumour-samples compared to matched non-transformed tissue samples (Figure 2I).

### Upregulation of USP10 is an early event in CRC and conserved in ex vivo human and genetically engineered murine models of colorectal cancer

Publicly available data did highlight that expression of *USP10* and *CTNNB1* were elevated irrespective of CRC stage (Figure S2F, G). We tested this observation in human and murine intestinal wild typic and tumour organoid models regarding the regulation of USP10 (Figure 2K-M). Similar to the observation in patent-derived primary resected CRC tumours, the endogenous protein levels of USP10 was significantly increased in tumour derived organoids, compared to non-oncogenic (Figure 2K). This was further supported by analysing publicly available expression data of patient derived non-oncogenic and CRC organoids (van de Wetering et al., 2015) (Figure 2L). Next, we wondered if genetic complexity could be a contributor to USP10 upregulation. To address this question, we utilised murine wild type organoids (*Cas9*) and employed CRISPR gene editing to delete/ mutate *Apc:Kras*^*G12D*^ (*AK*) or *Apc:Trp53:Kras*^*G12D*^ (*APK*), respectively (Figure 2M and S2H,I). Not only did loss of *Apc* induce morphologic changes of wild type murine organoids, but also alleviated the requirement for WNT activating components in the growth medium. This is in accordance with previous reports, and mutation of *Kras* to *Kras*^*G12D*^ upregulated endogenous Erk1/2 signalling and alleviated the requirement for EGF supplementation (Figure S2I) (Matano et al., 2015). Immunoblotting of Usp10 and β-Catenin confirmed that Usp10 was significantly enriched in transformed organoids when compared to parental control, and this was consistent with increased abundance of β-Catenin (Figure 2M).

These data demonstrate that the upregulation of USP10 in colorectal cancer is an early event, caused by oncogenic transformation irrespective of genetic driver complexity, and coincides with elevated abundance of the WNT effector β-Catenin.

### Truncation of APC allows for de novo protein-protein interaction between USP10 and β-Catenin in CRC

We hypothesized that β-Catenin directly interacts with USP10 in CRC and tested whether mutations within either β-Catenin or *APC* are a prerequisite to enable a protein-protein interaction. To this end, we co-immunoprecipitation endogenous USP10 and β-Catenin in the human CRC lines HCT116 (*CTNNB1*^*mutant*^*/APC*^*wildtype*^) and HT-29 (*CTNNB1*^*wildtype*^*/APC*^*truncated*^, Figure 3A, B). While USP10 and β-Catenin were singly immunoprecipitated in HCT116, no co-precipitation was observed. In contrast, USP10 co-immunoprecipitated with endogenous β-Catenin in HT-29 cells (Figure 3A). This observation highlighted the possibility that the truncation status and remaining length of APC is involved in the interaction between USP10 and β-Catenin. This was assessed by using a panel of human CRC lines, comprising LS174T, DLD-1, SW480, SW620, Caco-2 and Colo320, which harbour varying truncation mutations within *APC* (Figure S3A). Endogenous co-immunoprecipitation of USP10 and β-Catenin only occurred in Colo320, confirming that a proximal truncation in *APC* is a prerequisite, as cell lines carrying distal deletions, such as DLD-1, Caco-2 and SW480/SW620, failed to co-immunoprecipitate USP10 with β-Catenin (Figure S3B).

**Figure 3:**
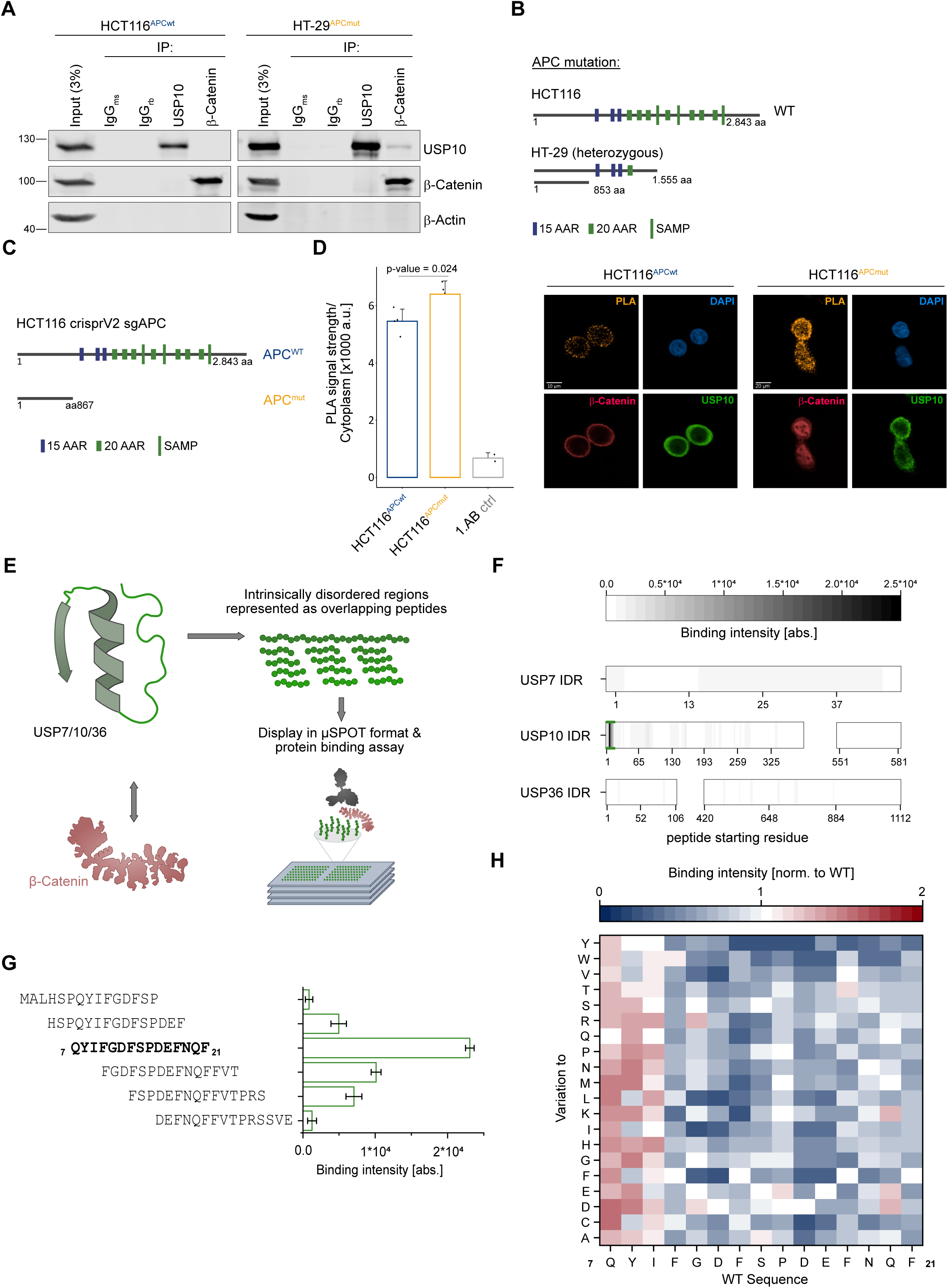
Truncation of APC allows for de novo protein-protein interaction between USP10 and β-Catenin in CRC. A. Representative input and endogenous co-immunoprecipitation of USP10 and β-Catenin in human CRC cell lines either wild type for APC, HCT116^APCwt^, or truncated HT-29^APCmut^. IgG served as antibody specificity control. β -Actin served as loading control. Input represents 3% of total loading. n=3. B. Schematic representation of truncating mutations reported in the *APC* gene in the CRC cell lines HCT116 and HT-29. Dark blue box = 15 AAR domains, green small boxes = 20 AAR domains, large green boxes = SAMP domains. 15- and 20-AAR = β-Catenin amino acid repeats; SAMP = Axin binding sites. Images adapted from the publicly available database www.uniprot.org. C. Schematic model of acute truncation of *APC* at amino acid 867 in HCT116 via CRISPR gene editing. D. Bargraph of PLA between USP10 and β-Catenin in either *APC wild type (APC*^*wt*^) or *APC*^*867*^ truncated (APC^mut^) HCT116. Data analysed from more than 750 cells over two independent experiments per condition. P-values were calculated using Mann-Whitney U test. Representative immunofluorescent images of endogenous USP10 (green), β-Catenin (red) and the corresponding PLA (mustard) in either *APC*^*wt*^ of *APC*^*mut*^ HCT116. DAPI served as nuclear marker. n=2. E. Schematic overview of the *in vitro* binding assay in µSPOT format. Intrinsically disordered regions of USP7, USP10 and USP36, respectively, were determined by <50 pLDDT score in their individual AlphaFold2 structural prediction and represented as 15 mer peptides overlapping 12 or 11 amino acids. For binding assays, µSPOT slides bearing the peptide library were incubated with recombinantly expressed and purified β-Catenin. β-Catenin binding to the on-chip peptides was detected by immunostaining with a chemiluminescent readout. F. Overview of binding intensities of recombinant β-Catenin towards discreet unstructured regions of USP7, USP10 and USP36. Gaps indicate the presence of structured domains within the DUBs. Colour code indicates binding intensity. Peptides with the globally highest binding intensity (N-terminal region of USP10 residues 7-21) are underlined in green and represented as a bar graph in panel G). Mean of n=3. G. Identified amino acid sequence within the N-terminal unstructured part of USP10 binding to recombinant β-Catenin. Bar graph shows binding intensity (abs. = absolute intensity). Mean of n=3 with corresponding standard deviation. H. Full positional scan of the most prominent β-Catenin binding hotspot USP10^7-21^ identified in the overlapping scan (panel G). Each residue of the peptide sequence was systematically varied to every other proteogenic amino acid and their β-Catenin binding intensities are shown relative to the wildtype sequence. Note that amino acid variations for certain positions result in drastic reductions in binding intensity compared to the wildtype sequence, thus suggesting direct interactions of the respective sidechains with β-Catenin. Mean of n=3.

Furthermore, based on the genetic alterations reported for HT-29 and Colo320, we concluded that the 15- and 20 β-Catenin binding amino acid repeat (AAR) domains within APC are required to directly compete for binding of USP10 to β-Catenin. Hence, we postulate that the putative de novo interaction between USP10 and β-Catenin requires the loss of the AAR domains. Proximity ligation assays (PLA) between USP10 and β-Catenin, carried out in CRISPR mediated truncation of *APC* in HCT116 cells confirmed that *APC* competed with USP10 for binding to β-Catenin (Figure 3C, D).

To confirm the interaction and map the USP10 binding site required for β-Catenin interaction, we conducted a µSPOT protein binding assay (Schulte et al., 2022). Here, the intrinsically disordered regions (IDR) of USP10, along with the IDR sequences of a known β-Catenin binder, USP7 (Novellasdemunt et al., 2017), and an additional DUB comprised of large unstructured regions, USP36 were displayed as overlapping peptide libraries and probed with recombinant β-Catenin (Figure 3E). We identified USP10 residues ^7^QYIFGDFSPDEFNQF^21^ (Figure 3F-H) to mediate direct and robust binding to β-Catenin. Intriguingly, AlphaFold2 MultimerV1.0 (AF2M) (Jumper et al., 2021), with the complete sequences as input, predicts the same residues within USP10 to engage with β-Catenin. Intriguingly, this binding site is overlapping with APC binding to β-Catenin (Figure S3C).

Taken together, this data confirms a direct USP10-β-Catenin interaction and reveals a competition of USP10 with APC for an identical β-Catenin binding site. Thus, lending a molecular explanation for the observed indirect β-Catenin stabilizing effect of APC truncations.

### Acute deletion of Usp10 in intestinal stem cells of D.melanogaster rescues hyperplasia and lethality of the Apc^Q8/Q8^ model

As USP10 is highly conserved between species we used *D*.*melanogaster* to investigate its involvement in intestinal homeostasis and hyperproliferation upon loss of function mutations within *APC*, (Martorell et al., 2014) (Figure 4 and S4). Firstly, we assessed the impact of Usp10 silencing on intestinal progenitor homeostasis. Intestinal progenitor cells were marked by GFP expression, driven under the control of the *escargot* regulatory region (*esg*::GAL>GFP), and immunofluorescence against armadillo, the *D*.*melanogaster* homologue to β-Catenin, that is expressed on the surface of intestinal stem cells (ISCs, Figure S4A). Expression of shRNA against Usp10 resulted in a marked reduction of ISCs when compared to a LacZ control shRNA (Figure S4A and B). Moreover, to establish a genetic linkage between *APC* truncation and USP10 in a tumour-like setting, we used the *Apc*^*Q8*^ hyperplasia model (McCartney et al., 2006). The allele *Apc*^*Q8*^ contains a premature stop codon leading to a significant truncation of *Apc* and loss of the β-catenin binding sites. Immunofluorescent analysis of *D*.*melanogaster* midguts revealed that heterozygous loss of *Apc* had a minor effect on overall tissue homeostasis highly similar to wildtype midguts. In contrast, midguts derived from adult animals carrying a homozygous *LOF truncating* mutation within *Apc* (*Apc*^*Q8/Q8*^) presented an entirely disorganised intestine (Figure 4A). This midgut was robustly populated by escargot-positive progenitors, many of them expressing the stem cell marker and Notch ligand Delta. (Figure 4A and B). Knockdown of Usp10, however, suppressed the stem cell and progenitor expansion observed in homozygous *Apc*^*Q8*^, and animals presented a midgut resembling a normal appearance (Figure 4A and B). Analysis of isolated midguts from either *Apc*^*Q8/+*^, *Apc*^*Q8/Q8*^ *or Apc*^*Q8/Q8*^ flies expressing an shRNA against Usp10 (*Usp10i;Apc*^*Q8/Q8*^) in intestinal stem cells indicated a significant reduction in overall Usp10 transcript abundance, along with reduced expression of the stem cell marker escargot (Figure 4C and S4E).

**Figure 4:**
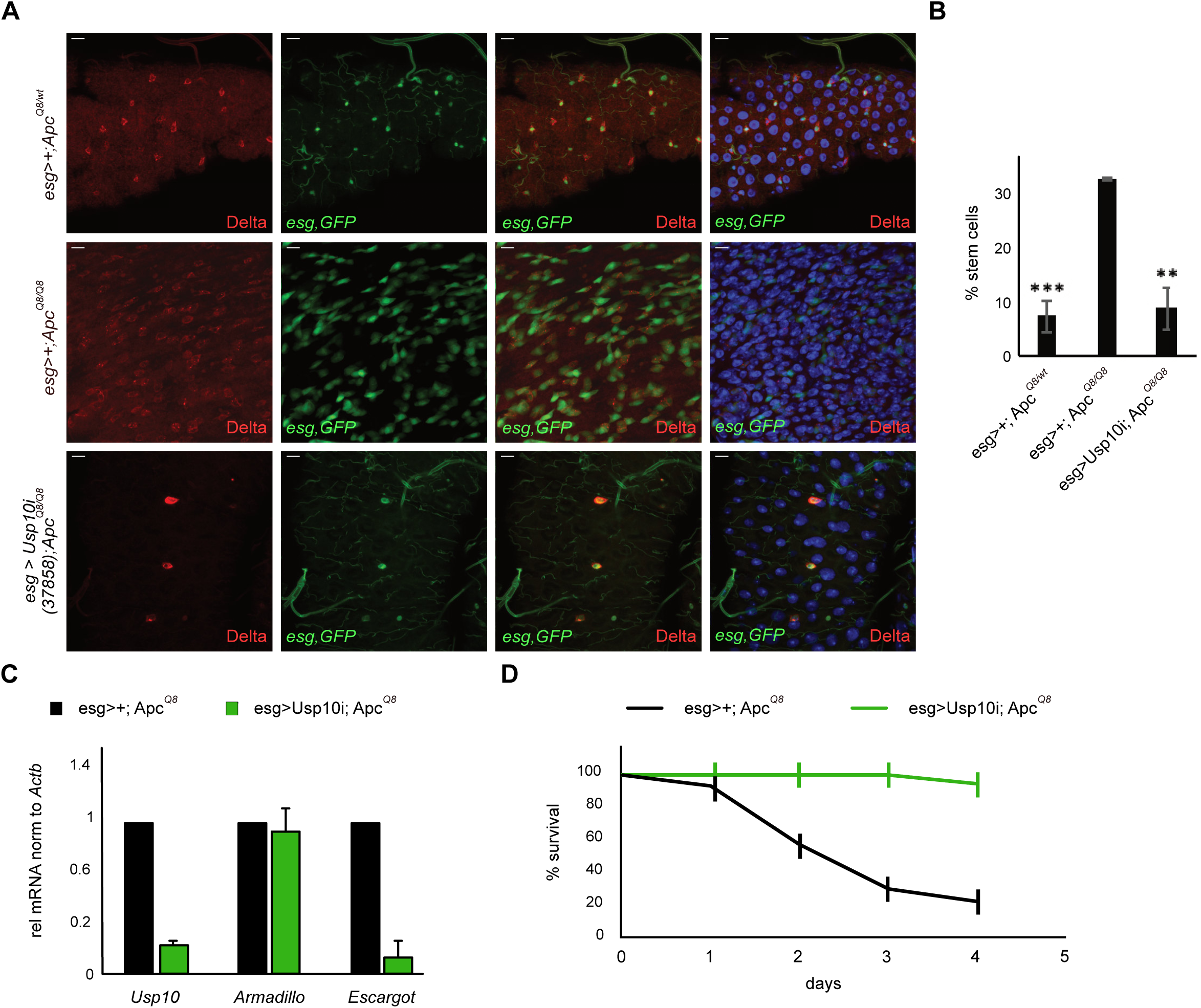
Acute deletion of Usp10 in intestinal stem cells of D.melanogaster rescues hyperplasia and lethality of the Apc^Q8/Q8^ mutant flies. A. Representative immunofluorescence of fly midguts. *Apc*^*Q8/+*^ heterozygotes are highly similar to wildtype midguts (not shown). Midguts of homozygous *Apc*^*Q8*^ mutants exhibit hyperproliferation of ISC (positive for the intestinal stem cell marker Delta (red)). Elimination of Usp10 using Usp10 inverted repeats (UAS-IR) suppresses the progenitor hyperproliferation phenotype observed in midguts of homozygous *Apc*^*Q8*^ mutants. B. Quantification of total stem cell abundance in all three conditions. Significance as compared to “esg>+; ApcQ8” was calculated using one-way ANOVA. **p<0.005; *** p<0.001. C. qRT-PCR analysis of the expression of *Usp10, armadillo* and *escargot* in midguts isolated from either *Apc*^*Q8*^ or *Apc*^*Q8*^ *Usp10*^*KD*^ flies. mRNA was normalised to *Actb*. Error bars represent standard deviation of 3 biological replicates. D. Kaplan Meier plot of adult survival of the indicated genotypes. *Apc*^*Q8*^ n=24, *Apc*^*Q8:esg-Usp10i*^ n=17.

Lastly, we investigated the impact of Usp10 deletion on overall survival in the background of *Apc*-truncation driven hyperplasia model. While the survival of heterozygous *Apc*^*Q8/wt*^ flies was similar to wildtype files, homozygous *Apc*^*Q8/Q8*^ mutation induced a temperature-sensitive lethality (Figure 4D and S4F). Strikingly, expression of an shRNA against *Usp10* in intestinal progenitors restored longevity, likely by negating the adverse effects on overall tissue homeostasis and growth imprinted by *Apc*^*Q8/Q8*^ (Figure 4D).

These data demonstrate an epistatic genetic linkage between USP10 and truncated APC that is required for ectopic stem cell proliferation.

### USP10, via controlling β-Catenin protein stability, regulates WNT signalling and stemness signature genes

To further elucidate the function of USP10 in CRC, we deleted endogenous USP10 by co-targeting of exon 2 and 10 in HT-29 and HCT116, respectively (Figure 5A and S5A, B). Depletion of USP10 in HT-29 resulted in a marked reduction of β-Catenin protein, but not its mRNA transcript, along with reduction in the CRC protein marker and WNT target gene LGR5 (Figure 5A and B). Loss of USP10 enhanced overall ubiquitylation of β-Catenin (Figure 5C) and accelerated protein turnover in HT-29 cells (Figure 5D and quantified in 5E). Depletion of USP10 in HCT116, however, had no effect on overall β-Catenin abundance nor ubiquitylation (Figure S5B-D), confirming the dependency of the USP10-β-Catenin interaction on *APC*-truncation. Interestingly, while cells deleted for USP10 by CRISPR mediated targeting did show reduced abundance and increased ubiquitylation of β-Catenin, longitudinal propagation of HT-29^ΔUSP10^ was not possible. Targeted cells within a heterogeneous cell pool were rapidly outcompeted by wildtype cells (Figure S5E). This is in line with previous reports of cell lethality upon USP10 loss (Higuchi et al., 2016). To by-pass this long-term lethality, we used an inducible knock down system, comprising two independent shRNA against *USP10*, to acutely deplete the DUB in HT-29 (Figure S5F). USP10 depleted HT-29 showed a significantly reduced proliferation, when compared to control vector transduced cells (Figure S5G).

**Figure 5:**
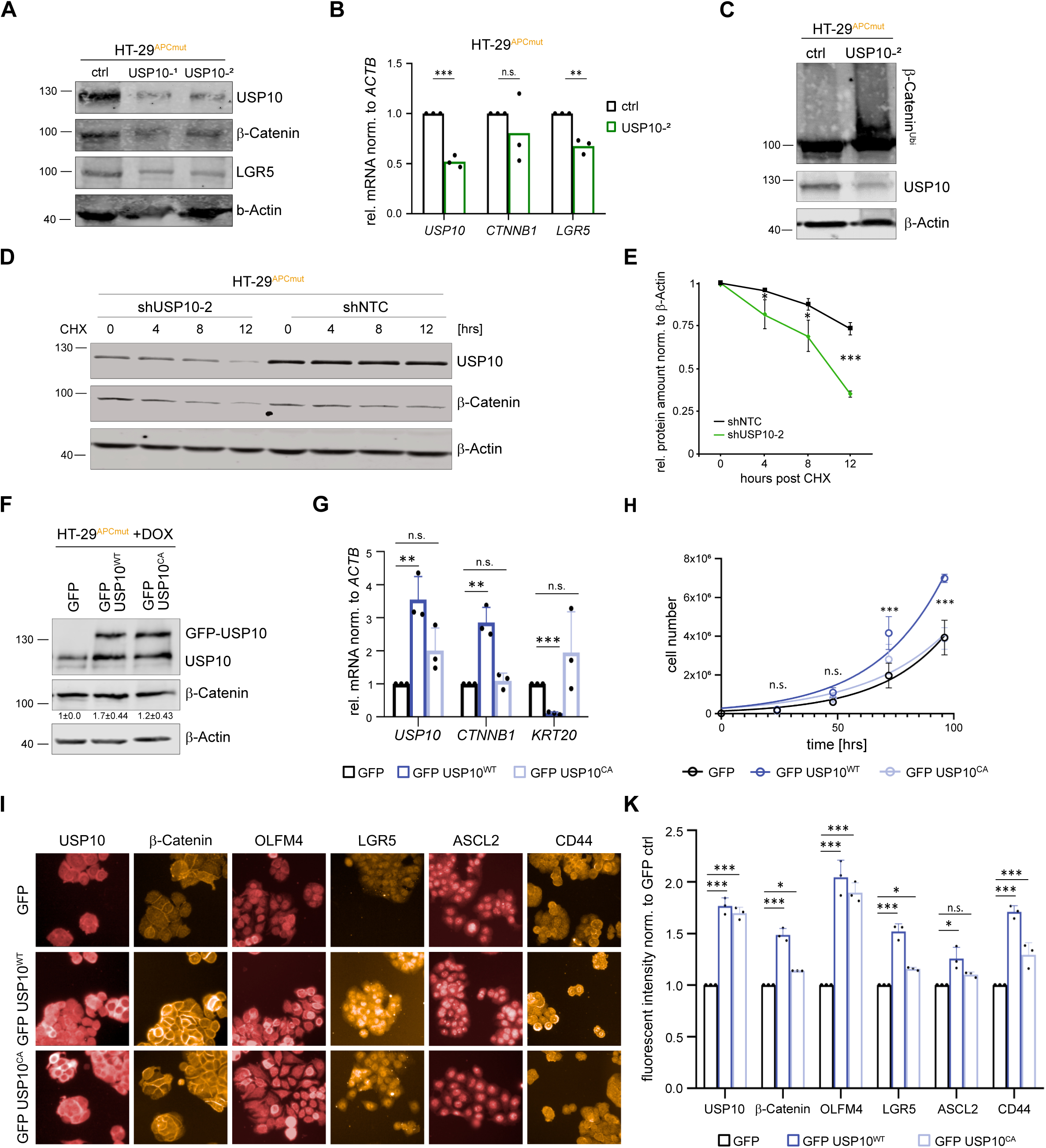
*USP10 regulates WNT signalling and stemness signature genes* via controlling β-Catenin protein stability. A. Immunoblot against endogenous USP10, β-Catenin and LGR5 in *APC* mutant HT-29 cells upon CRISPR mediated depletion of USP10. Two different cell pools (USP10^−1^ and USP10^−2^) along with non-targeting control (ctrl) cells are shown. β-Actin served as loading control. n=3. B. Quantitative RT-PCR of *USP10, CTNNB1* and *LGR5* expression of HT-29 USP10 CRISPR pool (USP10^−2^) compared to control (ctrl) cells. Error bars represent standard deviation of n=3 independent experiments. Significance was calculated using students t-test. **p< 0.005; *** p< 0.001. n.s. = non-significant. C. Tandem Ubiquitin Binding Entity (TUBE) assay of endogenous poly-ubiquitylated proteins, followed by immunoblotting against endogenous β-Catenin in HT-29 USP10 CRISPR cells (USP10^−2^). Immunoblot against endogenous USP10 is shown. β-Actin served as loading control. n=2. D. Cycloheximide (CHX) chase assay (100 μg/ml) of control (shNTC) or shUSP10-2 expressing HT-29 cells for indicated time points. Representative immunoblot analysis of USP10 and β-Catenin. β-Actin served as loading control. n=3 E. Quantification of relative protein abundance of β-Catenin, normalised to β-Actin, as shown in D. Significance was calculated using students t-test. n=3 *p< 0.05; *** p< 0.001 F. Representative immunoblot against endogenous USP10 and β-Catenin in *APC* mutant HT-29 cells upon DOX-inducible overexpression of GFP control (GFP), catalytical active GFP-USP10 (GFP USP10^WT^) and a catalytical inactive mutant of USP10 (GFP USP10^CA^). β-Actin served as loading control. (n=3) G. Quantitative RT-PCR of *USP10, CTNNB1* and *KRT20* expression of HT-29 cells overexpressing exogenous USP10. Error bars represent standard deviation of n=3 independent experiments. Significance was calculated using students t-test. ** p< 0.005; *** p< 0.001. n.s. = non-significant. H. Growth-curve of GFP USP10^WT^ and GFP USP10^CA^ overexpressing HT-29 cells compared to GFP control cells. Error bars represent standard deviation of n=3 independent experiments. Significance was calculated using one-way ANOVA. *** p< 0.001. n.s. = non-significant. I. Representative immunofluorescence images of conditional USP10^WT^ and USP10^CA^ overexpression and GFP control in HT-29 cells. K. Quantification of I. Mean intensity over well was measured and normalised to GFP control. Error bars represent standard deviation of n=3. Significance was calculated using unpaired t-test. *p<0.05; ***p<0.001; n.s. = non-significant.

A stem niche specific contribution of USP10 was further supported by analysing the whole proteome of HT-29 cells treated with either non-targeting (ctrl) or USP10 siRNA for 24 hours (Figure S5H, I). Among the downregulated proteins were proteins associated with the stem cell niche, including TCF4 (TCF7L2), TNFRSF21, NOTCH2, LGR4, CD44, along with reduced protein level of the proto-oncogene MYC, a direct target of WNT signalling (Figure S5H).

To investigate the extent of regulation of the WNT effector β-Catenin by USP10, and using a gain-of-function approach, we conditionally overexpressed either wild type (USP10^WT^) or a catalytic inactive variant of USP10 (USP10^CA^) in the CRC line HT-29. Conditional increase in USP10 led to an increase in β-Catenin abundance on protein as well as mRNA level (Figure 5F, G). The catalytic activity of USP10 is required to facilitate these effects on β-Catenin, as USP10^CA^ failed to stabilise β-Catenin (Figure 5F, G). Expression of USP10 significantly enhanced overall proliferation of HT-29 cells, when compared to vector or catalytic inactive mutant control cells (Figure 5H). Given that β-Catenin directly controls intestinal homeostasis and stem- and cancer cell maintenance, next, we wondered if USP10 affects the expression of essential stem- and CRC pathways. Immunofluorescence imaging of HT-29 expressing either USP10^WT^ or USP10^CA^ demonstrated that proteins associated with the CSC stem niche, such as β-Catenin, OLFM4, LGR5, ASCL2 or CD44 were significantly upregulated in a USP10^WT^ dependent fashion (Figure 5I, K).

These observations establish that USP10 regulates the ubiquitylation and abundance of β-Catenin in an *APC* truncation dependent manner, thereby controlling WNT signalling and affecting the expression of (cancer) stem cell signature genes and CRC growth.

### USP10 is required to maintain CRC cell identity, stemness and tumour growth

To elucidate the involvement of Usp10 in the maintenance of established CRC, we utilized murine intestinal organoid cultures (Figure 2M). *APK* organoids were transduced with AAV encoding either shRNA against Usp10 or non-targeting control, respectively (Figure 6A, B and S6A). Transcriptomic analysis of *APK* organoids revealed that Usp10 is required to maintain WNT signalling, as loss of Usp10 significantly reduced this pathway (Figure 6C-E). Not only did the knock down of Usp10 result in reduced abundance of β-Catenin, but the expression of the WNT signalling target gene Myc and Ccnd2 were reduced, too (Figure 6E and S6C). Overall, reduction of Usp10 in *APK* organoids reduced signatures associated with stemness and induced the expression of differentiation gene signatures (Figure 6C-E and S6B, C).

**Figure 6:**
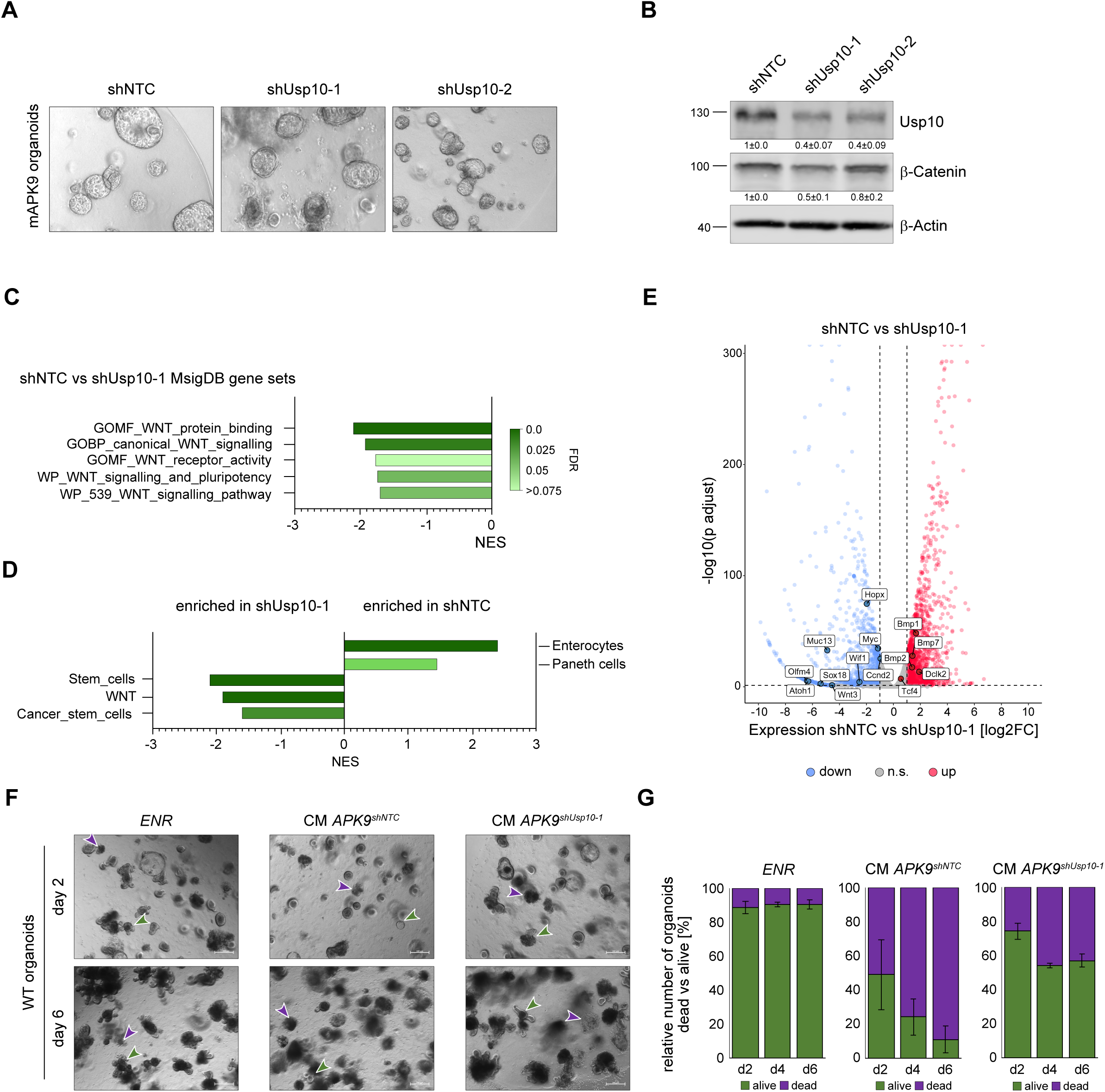
USP10 is required to maintain CRC cell identity, stemness and tumour growth. A. Representative brightfield images of stable transformed murine *A*^*ex9*^*PK*^*G12D*^ organoids (APK9). Two different shRNAs against Usp10 and shNTC expressing organoids were generated. B. Representative immunoblot of Usp10 and β-Catenin protein upon shRNA mediated knock-down of endogenous Usp10. β-Actin served as loading control. Quantification was calculated from n=3. C. Gene set enrichment analysis of MsigDB gene sets, deregulated in shUsp10-1 compared to shNTC APK9 organoids. D. Gene set enrichment analysis of intestinal specific gene sets, deregulated in shUsp10-1 compared to shNTC APK9 organoids. E. Volcano-plot of differential expressed genes upon knock-down of Usp10 in APK9 organoids. Up- and down-regulated genes are highlighted in red and blue, respectively. Genes-of-interest are labelled. F. Representative brightfield images of wild-type (WT) organoids cultured in ENR medium, *APK*^*shNTC*^ *and APK*^*shUsp10-1*^ conditioned medium (CM), supplemented with EGF and R-spondin, for up to 6 days. Purple arrows indicate dead organoids, green arrows indicate living organoids. E – EGF, N – Noggin, R - R-spondin. n=3. G. Quantification of F. Dead and alive organoids were counted and bar graphs represent percentage of alive vs dead organoids. Error bars represent standard deviation calculated from n=3 independent experiments.

These results show that USP10, in CRC at least, contributes to the control of differentiation and can be linked to intestinal cancer cell identity. Hence, USP10, via β-Catenin, promotes intestinal cancer stemness and propagation.

### Loss of Usp10 opposes competitor signalling and restores a wild-typic niche

Recently, a crosstalk between colorectal cancer and non-transformed intestinal stem cells was shown to be mediated by clonal competition and super-competitor *Apc*-dependent *Notum* signalling, that induces the death of the naïve stem cells. This impact of the cancer cell on non-transformed neighbouring naive intestinal stem cells was shown to be crucial for tumour development (Flanagan et al., 2021a; Schmitt et al., 2022) (van Neerven et al., 2021). Given the strong impact on WNT signalling and the extended control of β-Catenin by USP10, we examined whether Usp10 is required for the super-competitor phenotype and, specifically, if silencing of USP10 could oppose this signalling axis. Towards this end, we cultured wild type organoids in the presence of pre-conditioned medium from either *APK*^*shNTC*^ or *APK*^*shUsp10*^ organoids and assessed wild type organoid survival (Figure 6F and S6D). While established wild type organoids grew in ENR medium, exposure to *APK*^*shNTC*^ derived medium rapidly resulted in wild type organoid loss (Figure 6F, G). Remarkably, when cultured in medium from *APK*^*shUsp10*^ organoids, most wild type organoids survived an extended culture under these conditions (Figure 6F, G and S6D). Analysis of the transcriptome of *APK*^*shUsp10-2*^ organoids revealed that *Notum*, along with genes associated with a super competitor signature, such as *Dkk2, Dkk3* or *Wif1* (Flanagan et al., 2021b; van Neerven et al., 2021), were downregulated upon loss of Usp10 (Figure S6C).

## Discussion

Colorectal cancer is mostly driven by deregulated WNT signalling, resulting in hyper-activation of the proto-oncoprotein β-Catenin and constant transcription of its target genes (Zhan et al., 2017). This is predominantly initiated via loss of function mutations within the tumour suppressor gene *APC*; around 80% of all CRC patients harbour these mutations (Gao et al., 2013; Siegel et al., 2021). It is noteworthy that truncation mutations vary regarding localisation and the therefore resulting remaining domains within the APC protein (Ranes et al., 2021). Recent studies have reported that tumour progression, loss-of-heterozygosity, and deregulation of WNT signalling are indeed affected by the length of the remaining tumour suppressor (Albuquerque et al., 2002). Furthermore, aberrant WNT signalling, caused by *APC* truncation, is a prerequisite for the establishing of a super competitor cell, manifesting CRC onset (Flanagan et al., 2021b; Schulte et al., 2022; van Neerven et al., 2021). Despite our growing understanding of the underlying genetic causalities of colorectal cancer, the identification of mechanisms at the basis of this phenomenon and identification of suitable therapeutic targets presents a challenge that yet needs to be overcome (Guren, 2019).

One promising strategy is targeting deregulated protein stability in cancer, in particular the Ubiquitin Proteasome System (UPS). Despite the loss of either APC or mutations within the degron motive of β-Catenin, the WNT effector is still ubiquitylated, at least, in CRC-derived cell lines (Ranes et al., 2021; Yang et al., 2006). Several E3 ligases have been reported in the past to ubiquitylate and thereby regulate the stability and activity of this proto-oncogene (Chitalia et al., 2008; Dominguez-Brauer et al., 2017; Jiang et al., 2016; Shearer et al., 2015; Thomas et al., 2016). This opens an intriguing possibility to target the protein levels of β-Catenin by interfering with enzymes conferring stabilisation. As a potential therapeutically relevant druggable family of proteins DUBs are of interest, as this class of enzymes opposes substrate ubiquitylation and can contribute to protein stabilisation and activation (Clague et al., 2019).

In this study, we investigated the possibility to control the abundance of β-Catenin protein via deubiquitylases in colorectal cancer driven by loss of the tumour suppressor APC. By siRNA interference screening in the APC truncated human CRC line HT-29, the deubiquitylase USP10 was identified as a novel putative regulator of β-Catenin abundance. Previously, USP10 was reported to control the protein stability of TP53 (Li et al., 2022; Yuan et al., 2010) and contribute to control of autophagy (Jia and Bonifacino, 2021; Liu et al., 2011), DNA damage (Bhattacharya et al., 2020; Zhang et al., 2016) and metabolic signalling (Bhattacharya et al., 2020; Deng et al., 2016), processes tumour cells highly rely on (Hanahan, 2022). Analysing publicly available patient data and samples from local CRC patients, we found that USP10 is frequently upregulated in human CRC tumours and is often co-expressed with β-Catenin. Intriguingly, by utilising a limited panel of human CRC cell lines, we observed that USP10 only interacted with β-Catenin when the truncation mutation within APC resulted in loss of the AAR domains. This observation was further corroborated by using CRISPR targeting of *APC* in otherwise *APC* non-mutant CRC lines. Microarray-based binding assays identified residues 7-21 of the unstructured N-terminus of USP10 to directly interact with β-Catenin. Comparison of the resulting structural model with the APC-β-Catenin complex lends a molecular explanation for the observed direct competition with APC. An overlapping binding site of APC and USP10 on β-Catenin would also rationalize the need for APC truncation for successful USP10 co-immunoprecipitation. Additionally, this observation could explain how APC mutations indirectly control β-Catenin abundance without affecting the ubiquitylation level *per se*. Taken together, our data extends the role of WNT signalling and β-Catenin in CRC and proposes patient stratification towards USP10 dependency.

While USP10 regulates the abundance of β-Catenin by affecting its ubiquitylation, its loss had widespread consequences for *APC* truncated CRC. Altering USP10 abundance affected the expression of genes associated with proliferation, stemness and disease progression. Usp10 imprints the cellular identity of cancer cells, as loss of Usp10 altered the transcriptional profiles towards a non-transformed, differentiated state. Not only does USP10 control tumour intrinsic pathways and biological processes regulated by WNT, loss of USP10 suppressed cell death of non-transformed cells exposed to cultured medium from tumour organoids. Given that the super competitor signalling cascade (Flanagan et al., 2021b; Schmitt et al., 2022; van Neerven et al., 2021) leads to the secretion of signalling molecules initiating cell death, loss of USP10 controls extrinsic signalling mechanisms and thereby opposes tumour growth. It is worth noting that the catalytic activity of USP10 was required for the effects observed; overexpression of the catalytic inactive form had no effect on global β-Catenin abundance nor on proliferation. Hence, the catalytic active site presents a suitable target site (Lange et al., 2022).

The observed effects in gene expression in the USP10 shRNA cell lines and organoids were a direct consequence of reduced β-Catenin protein abundance. These processes are highly conserved, as by employing *D*.*melanogaster* intestinal hyperproliferation models and silencing of *Usp10* in the intestinal stem cell niche demonstrated that targeting USP10 *in vivo*, indeed, did ameliorate the *APC* phenotype caused by homozygous loss of the tumour suppressor (*APC*^*Q8/Q8*^) (Martorell et al., 2014).

Utilizing human primary patient material, colorectal cancer cell lines as well as genetically tailored murine organoids and *Drosophila melanogaster* as a model system for aberrant WNT signalling in the intestine, we unravelled a novel protein-protein interaction to which CRC are addicted. Taken together, our study provided robust *in vitro* and *in vivo* evidence that USP10 functions as a driver of CRC and could serve as a therapeutic relevant target in a distinct subset of *APC*-truncated patient cohort.

## Supporting information

Reissland et al Suppl Figures 1-6

Reissland et al Material and Methods

## Ethics approval and Consent to participate

*In vivo* experiments concerning CRISPR mediated oncogenesis or the *Apc*^*min/+*^ mouse model were approved by the Regierung Unterfranken and the ethics committee under the license numbers 2532-2-555, 2532-2-556, 2532-2-694 and 2532-2-1002. The mouse strains used for this publication are listed. All animals are housed in standard cages in pathogen -free facilities on a 12 -h light/dark cycle with *ad libitum* access to food and water. FELASA2014 guidelines were followed for animal maintenance.

Mouse experiments concerning AOM/DSS induced oncogenesis were reviewed and approved by the Regierungspräsidium Darmstadt, Darmstadt, Germany.

Human colorectal cancer samples, irrespective of sex, were obtained from the Pathology Department at the University Hospital Würzburg (Germany). Informed consent was obtained from all patients. Experiments agreed with the principles set out in the WMA Declaration of Helsinki and the Department of Health and Human Services Belmont Report. Samples were approved under Ethics Approval 17/01/2006 (University Hospital Würzburg).

## Consent for publication

We have obtained consent to publish this paper from all the study participants.

## Author contributions

Conceptualization: M.R., M.E.D.

Methodology: M.R. (*in vitro*, Co-IP, *in vivo*, TUBE, organoids, RNA Seq., Bioinformatic); O.H. (*in vivo* IHC and IF), S.T. (*D. melanogaster*), C.P.G. & A.C.J. (Mass Spec.), D.S. & S.L. (PLA); C.F. (*in vitro*), N.P. (*in vitro*), M.P. & F.G. (in vivo transplant), J.B. & M.M. (Bioinformatic), C.S. & H.M. (SPOT Array), C.S.V. (Operetta system), C.A. (loading Ilumina), V.B. & M.Ro. (human FFPE samples), A.W. (resection material for human organoids).

Formal analysis: M.R. (*in vitro, in vivo*); M.R., J.B. (Bioinformatic), O.H., M.Ro. and M.E.D. (Pathology); S.T., A.O. & P.G. (*D*.*melanogaster*); C.S. & H.M. (SPOT); C.P.G.& I.D. (Mass Spec), M.P. & F.G. (in vivo) Investigation: M.R., O.H., H.M., A.O. M.Ro., P.G., M.E.D.

Resources: A.W., M.P., V.L., M.M., M.Ro., H.M., F.G., A.O., P.G., M.E.D.

Writing-original draft: M.E.D.

Writing-review and editing: M.R., V.L., A.W., M.M., M.E., H.M., F.G., M.Ro., A.O., P.G. M.E.D.

Supervision: M.E.D. Funding acquisition: M.E.D.

## Conflict of Interest

The authors declare no potential conflicts of interest.

## Acknowledgements

The Operetta High Content Microscope was funded by the Deutsche Forschungsgemeinschaft (DFG, German Research Foundation) -440766788 (INST 93/1023-1 -FUGG).

## Financial Support

MR is funded by the DKH MSNZ Wuerzburg, DFG-GRK 2243 and IZKF B335. OH is supported by the German Cancer Aid via grant 70112491 and 70114554. M.E. is supported by the TransOnc priority programme of the German Cancer Aid within grant 70112951 (ENABLE). AO and MED are funded by the German Israeli Foundation grant 1431 1431 and ICRF project grant to AO. Open Access funding enabled and organized by Projekt DEAL. H.M.M and C.S. are funded by the DFG (DFG MA6957/1-1).

---

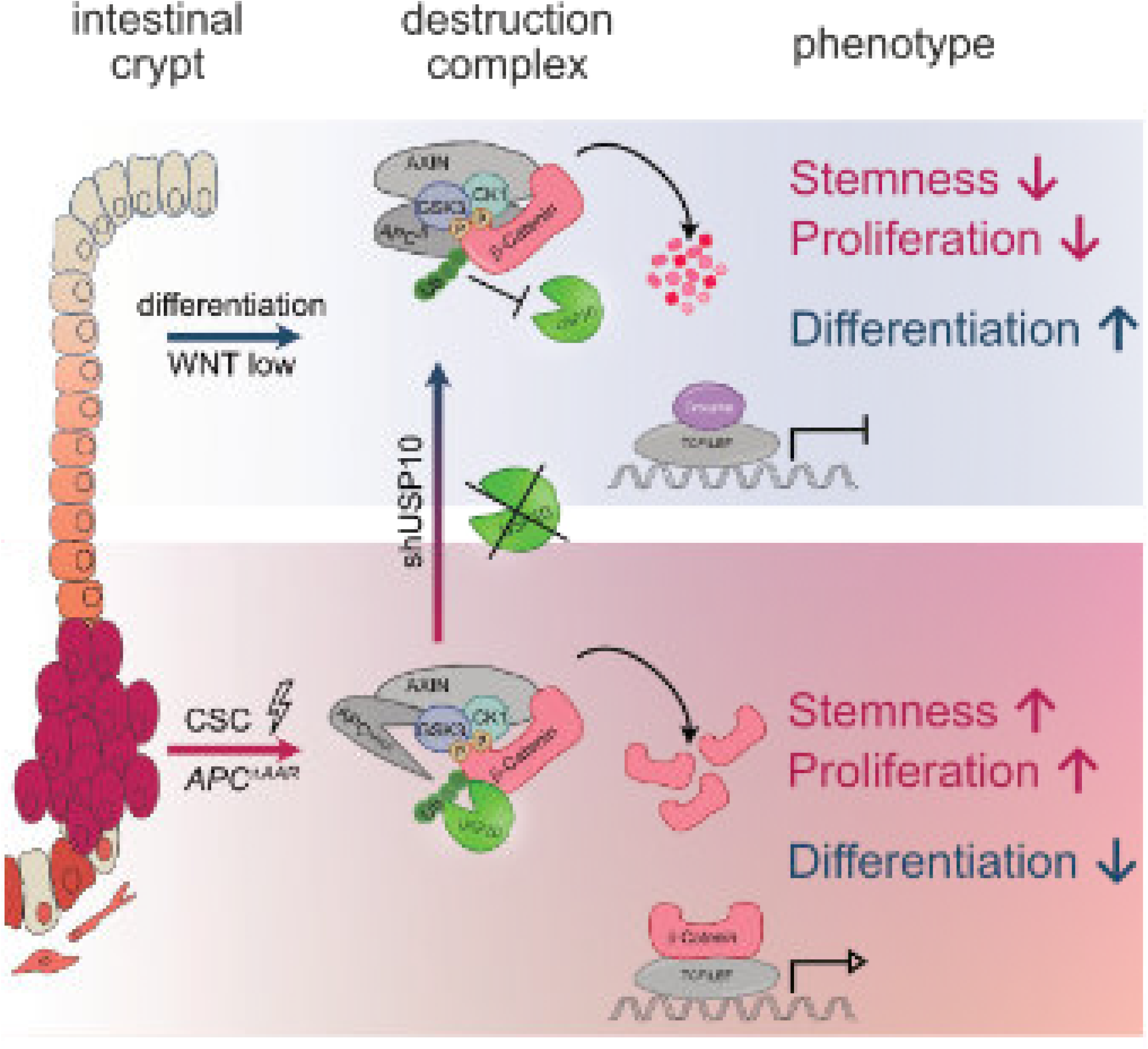

## Notes

### Competing Interest Statement

The authors have declared no competing interest.

### Summary of Updates

minor corrections

## Literature

.Risks and causes of bowel cancer | Cancer Research UK.

Ahmed, Y., Hayashi, S., Levine, A., and Wieschaus, E. (1998). Regulation of armadillo by a Drosophila APC inhibits neuronal apoptosis during retinal development. Cell 93, 1171–1182.

Albuquerque, C., Breukel, C., van der Luijt, R., Fidalgo, P., Lage, P., Slors, F.J., Leitao, C.N., Fodde, R., and Smits, R. (2002). The ‘just-right’ signaling model: APC somatic mutations are selected based on a specific level of activation of the beta-catenin signaling cascade. Hum Mol Genet 11, 1549–1560.

Barua, D., and Hlavacek, W.S. (2013). Modeling the effect of APC truncation on destruction complex function in colorectal cancer cells. PLoS Comput Biol 9, e1003217.

Bhattacharya, U., Neizer-Ashun, F., Mukherjee, P., and Bhattacharya, R. (2020). When the chains do not break: the role of USP10 in physiology and pathology. Cell Death Dis 11, 1033.

Cerami, E., Gao, J., Dogrusoz, U., Gross, B.E., Sumer, S.O., Aksoy, B.A., Jacobsen, A., Byrne, C.J., Heuer, M.L., Larsson, E., et al. (2012). The cBio cancer genomics portal: an open platform for exploring multidimensional cancer genomics data. Cancer Discov 2, 401–404.

Chen, C., Zhu, D., Zhang, H., Han, C., Xue, G., Zhu, T., Luo, J., and Kong, L. (2018). YAP-dependent ubiquitination and degradation of beta-catenin mediates inhibition of Wnt signalling induced by Physalin F in colorectal cancer. Cell Death Dis 9, 591.

Chitalia, V.C., Foy, R.L., Bachschmid, M.M., Zeng, L., Panchenko, M.V., Zhou, M.I., Bharti, A., Seldin, D.C., Lecker, S.H., Dominguez, I., et al. (2008). Jade-1 inhibits Wnt signalling by ubiquitylating beta-catenin and mediates Wnt pathway inhibition by pVHL. Nat Cell Biol 10, 1208–1216.

Clague, M.J., Urbe, S., and Komander, D. (2019). Breaking the chains: deubiquitylating enzyme specificity begets function. Nat Rev Mol Cell Biol 20, 338–352.

Deng, M., Yang, X., Qin, B., Liu, T., Zhang, H., Guo, W., Lee, S.B., Kim, J.J., Yuan, J., Pei, H., et al. (2016). Deubiquitination and Activation of AMPK by USP10. Mol Cell 61, 614–624.

Dominguez-Brauer, C., Khatun, R., Elia, A.J., Thu, K.L., Ramachandran, P., Baniasadi, S.P., Hao, Z., Jones, L.D., Haight, J., Sheng, Y., et al. (2017). E3 ubiquitin ligase Mule targets beta-catenin under conditions of hyperactive Wnt signaling. Proc Natl Acad Sci U S A 114, E1148–E1157.

Evans, R., O’Neill, M., Pritzel, A., Antropova, N., Senior, A., Green, T., Žídek, A., Bates, R., Blackwell, S., Yim, J., et al. (2021). Protein complex prediction with AlphaFold-Multimer. bioRxiv, 2021.2010.2004.463034.

Flanagan, D.J., Pentinmikko, N., Luopajarvi, K., Willis, N.J., Gilroy, K., Raven, A.P., McGarry, L., Englund, J.I., Webb, A.T., Scharaw, S., et al. (2021a). NOTUM from Apc-mutant cells biases clonal competition to initiate cancer. Nature 594, 430–435.

Flanagan, D.J., Pentinmikko, N., Luopajärvi, K., Willis, N.J., Gilroy, K., Raven, A.P., McGarry, L., Englund, J.I., Webb, A.T., Scharaw, S., et al. (2021b). NOTUM from Apc-mutant cells biases clonal competition to initiate cancer. Nature 2021 594:7863 594, 430–435.

Gao, C., Xiao, G., and Hu, J. (2014). Regulation of Wnt/beta-catenin signaling by posttranslational modifications. Cell Biosci 4, 13.

Gao, J., Aksoy, B.A., Dogrusoz, U., Dresdner, G., Gross, B., Sumer, S.O., Sun, Y., Jacobsen, A., Sinha, R., Larsson, E., et al. (2013). Integrative analysis of complex cancer genomics and clinical profiles using the cBioPortal. Sci Signal 6, pl1.

Guren, M.G. (2019). The global challenge of colorectal cancer. Lancet Gastroenterol Hepatol 4, 894–895.

Half, E., Bercovich, D., and Rozen, P. (2009). Familial adenomatous polyposis. Orphanet J Rare Dis 4, 22.

Hanahan, D. (2022). Hallmarks of Cancer: New Dimensions. Cancer Discov 12, 31–46.

Hankey, W., Frankel, W.L., and Groden, J. (2018). Functions of the APC tumor suppressor protein dependent and independent of canonical WNT signaling: implications for therapeutic targeting. Cancer Metastasis Rev 37, 159–172.

Higuchi, M., Kawamura, H., Matsuki, H., Hara, T., Takahashi, M., Saito, S., Saito, K., Jiang, S., Naito, M., Kiyonari, H., et al. (2016). USP10 Is an Essential Deubiquitinase for Hematopoiesis and Inhibits Apoptosis of Long-Term Hematopoietic Stem Cells. Stem Cell Reports 7, 1116–1129.

Jia, R., and Bonifacino, J.S. (2021). The ubiquitin isopeptidase USP10 deubiquitinates LC3B to increase LC3B levels and autophagic activity. J Biol Chem 296, 100405.

Jiang, J.X., Sun, C.Y., Tian, S., Yu, C., Chen, M.Y., and Zhang, H. (2016). Tumor suppressor Fbxw7 antagonizes WNT signaling by targeting beta-catenin for degradation in pancreatic cancer. Tumour Biol 37, 13893–13902.

Jumper, J., Evans, R., Pritzel, A., Green, T., Figurnov, M., Ronneberger, O., Tunyasuvunakool, K., Bates, R., Zidek, A., Potapenko, A., et al. (2021). Highly accurate protein structure prediction with AlphaFold. Nature 596, 583–589.

Lange, S.M., Armstrong, L.A., and Kulathu, Y. (2022). Deubiquitinases: From mechanisms to their inhibition by small molecules. Mol Cell 82, 15–29.

Li, H., Li, C., Zhai, W., Zhang, X., Li, L., Wu, B., Yu, B., Zhang, P., Li, J., Cui, C.P., et al. (2022). Destabilization of TP53 by USP10 is essential for neonatal autophagy and survival. Cell Rep 41, 111435.

Liu, J., Xia, H., Kim, M., Xu, L., Li, Y., Zhang, L., Cai, Y., Norberg, H.V., Zhang, T., Furuya, T., et al. (2011). Beclin1 controls the levels of p53 by regulating the deubiquitination activity of USP10 and USP13. Cell 147, 223–234.

Martorell, O., Merlos-Suarez, A., Campbell, K., Barriga, F.M., Christov, C.P., Miguel-Aliaga, I., Batlle, E., Casanova, J., and Casali, A. (2014). Conserved mechanisms of tumorigenesis in the Drosophila adult midgut. PLoS One 9, e88413.

Matano, M., Date, S., Shimokawa, M., Takano, A., Fujii, M., Ohta, Y., Watanabe, T., Kanai, T., and Sato, T. (2015). Modeling colorectal cancer using CRISPR-Cas9-mediated engineering of human intestinal organoids. Nat Med 21, 256–262.

McCartney, B.M., Price, M.H., Webb, R.L., Hayden, M.A., Holot, L.M., Zhou, M., Bejsovec, A., and Peifer, M. (2006). Testing hypotheses for the functions of APC family proteins using null and truncation alleles in Drosophila. Development 133, 2407–2418.

Morin, P.J., Sparks, A.B., Korinek, V., Barker, N., Clevers, H., Vogelstein, B., and Kinzler, K.W. (1997). Activation of beta-catenin-Tcf signaling in colon cancer by mutations in beta-catenin or APC. Science 275, 1787–1790.

Novellasdemunt, L., Antas, P., and Li, V.S. (2015). Targeting Wnt signaling in colorectal cancer. A Review in the Theme: Cell Signaling: Proteins, Pathways and Mechanisms. Am J Physiol Cell Physiol 309, C511–521.

Novellasdemunt, L., Foglizzo, V., Cuadrado, L., Antas, P., Kucharska, A., Encheva, V., Snijders, A.P., and Li, V.S.W. (2017). USP7 Is a Tumor-Specific WNT Activator for APC-Mutated Colorectal Cancer by Mediating beta-Catenin Deubiquitination. Cell Rep 21, 612–627.

Park, H.B., Kim, J.W., and Baek, K.H. (2020). Regulation of Wnt Signaling through Ubiquitination and Deubiquitination in Cancers. Int J Mol Sci 21.

Ranes, M., Zaleska, M., Sakalas, S., Knight, R., and Guettler, S. (2021). Reconstitution of the destruction complex defines roles of AXIN polymers and APC in beta-catenin capture, phosphorylation, and ubiquitylation. Mol Cell 81, 3246–3261 e3211.

Schmitt, M., Ceteci, F., Gupta, J., Pesic, M., Bottger, T.W., Nicolas, A.M., Kennel, K.B., Engel, E., Schewe, M., Callak Kirisozu, A., et al. (2022). Colon tumour cell death causes mTOR dependence by paracrine P2X4 stimulation. Nature.

Schneikert, J., Grohmann, A., and Behrens, J. (2007). Truncated APC regulates the transcriptional activity of beta-catenin in a cell cycle dependent manner. Hum Mol Genet 16, 199–209.

Schulte, C., Solda, A., Spanig, S., Adams, N., Bekic, I., Streicher, W., Heider, D., Strasser, R., and Maric, H.M. (2022). Multivalent binding kinetics resolved by fluorescence proximity sensing. Commun Biol 5, 1070.

Shearer, R.F., Iconomou, M., Watts, C.K., and Saunders, D.N. (2015). Functional Roles of the E3 Ubiquitin Ligase UBR5 in Cancer. Mol Cancer Res 13, 1523–1532.

Siegel, R.L., Miller, K.D., Fuchs, H.E., and Jemal, A. (2021). Cancer Statistics, 2021. CA: A Cancer Journal for Clinicians 71, 7–33.

Sung, H., Ferlay, J., Siegel, R.L., Laversanne, M., Soerjomataram, I., Jemal, A., and Bray, F. (2021). Global Cancer Statistics 2020: GLOBOCAN Estimates of Incidence and Mortality Worldwide for 36 Cancers in 185 Countries. CA Cancer J Clin 71, 209–249.

Thomas, J.J., Abed, M., Heuberger, J., Novak, R., Zohar, Y., Beltran Lopez, A.P., Trausch-Azar, J.S., Ilagan, M.X.G., Benhamou, D., Dittmar, G., et al. (2016). RNF4-Dependent Oncogene Activation by Protein Stabilization. Cell Rep 16, 3388–3400.

van de Wetering, M., Francies, H.E., Francis, J.M., Bounova, G., Iorio, F., Pronk, A., van Houdt, W., van Gorp, J., Taylor-Weiner, A., Kester, L., et al. (2015). Prospective derivation of a living organoid biobank of colorectal cancer patients. Cell 161, 933–945.

van Neerven, S.M., de Groot, N.E., Nijman, L.E., Scicluna, B.P., van Driel, M.S., Lecca, M.C., Warmerdam, D.O., Kakkar, V., Moreno, L.F., Vieira Braga, F.A., et al. (2021). Apc-mutant cells act as supercompetitors in intestinal tumour initiation. Nature 594, 436–441.

Wu, C., Luo, K., Zhao, F., Yin, P., Song, Y., Deng, M., Huang, J., Chen, Y., Li, L., Lee, S., et al. (2018). USP20 positively regulates tumorigenesis and chemoresistance through beta-catenin stabilization. Cell Death Differ 25, 1855–1869.

Yang, J., Zhang, W., Evans, P.M., Chen, X., He, X., and Liu, C. (2006). Adenomatous polyposis coli (APC) differentially regulates beta-catenin phosphorylation and ubiquitination in colon cancer cells. J Biol Chem 281, 17751–17757.

Yuan, J., Luo, K., Zhang, L., Cheville, J.C., and Lou, Z. (2010). USP10 regulates p53 localization and stability by deubiquitinating p53. Cell 140, 384–396.

Zhan, T., Rindtorff, N., and Boutros, M. (2017). Wnt signaling in cancer. Oncogene 36, 1461–1473.

Zhang, M., Hu, C., Tong, D., Xiang, S., Williams, K., Bai, W., Li, G.M., Bepler, G., and Zhang, X. (2016). Ubiquitin-specific Peptidase 10 (USP10) Deubiquitinates and Stabilizes MutS Homolog 2 (MSH2) to Regulate Cellular Sensitivity to DNA Damage. J Biol Chem 291, 10783–10791.

Zhao, H., Ming, T., Tang, S., Ren, S., Yang, H., Liu, M., Tao, Q., and Xu, H. (2022). Wnt signaling in colorectal cancer: pathogenic role and therapeutic target. Mol Cancer 21, 144.

