## Supplementary material for "Stabilisation of β-Catenin-WNT signalling by USP10 in APC-*truncated* colorectal cancer drives cancer stemness and enables super-competitor signalling": Reissland et al Suppl Figures 1-6

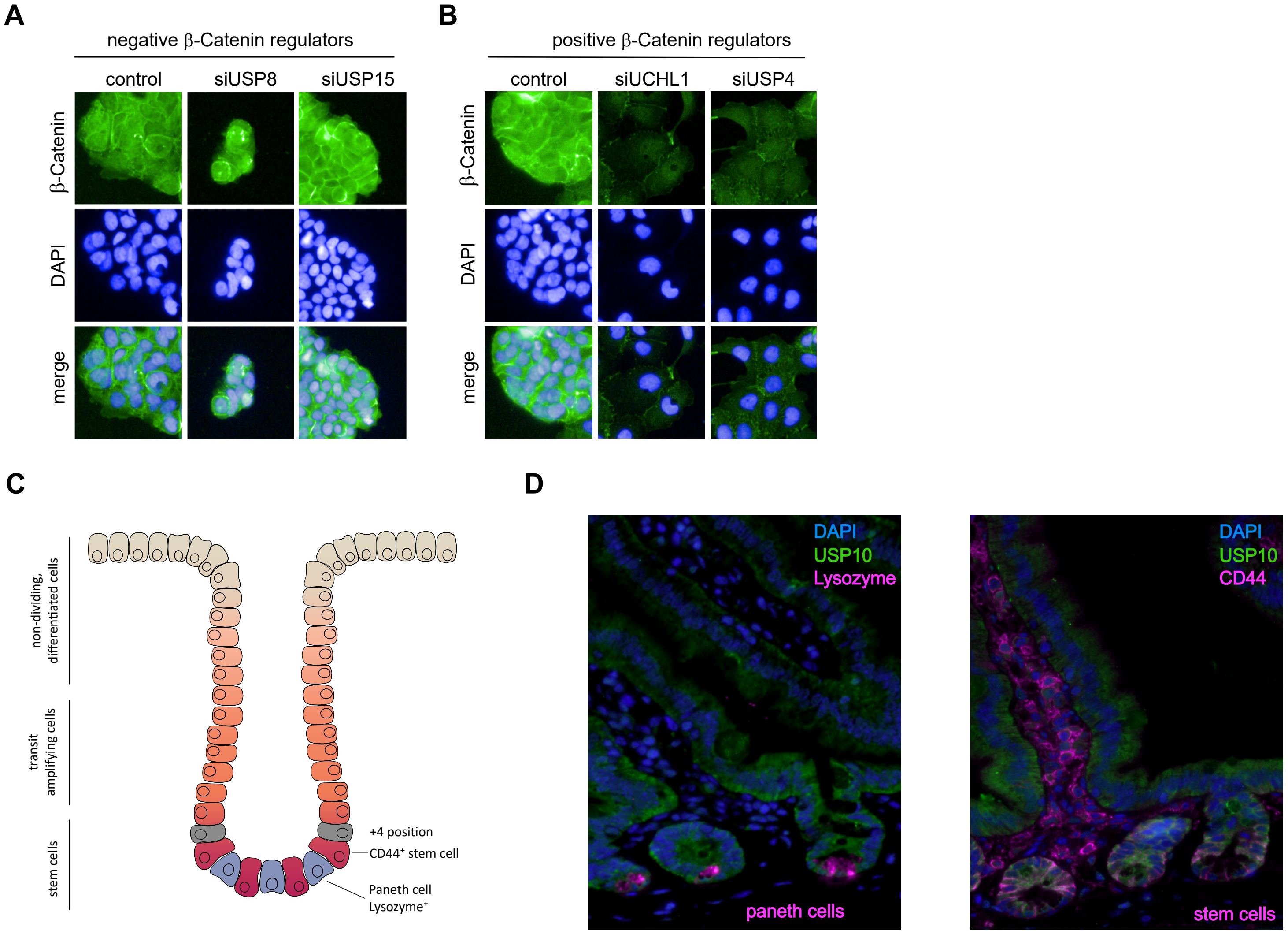


*Supplementary Figure 1*

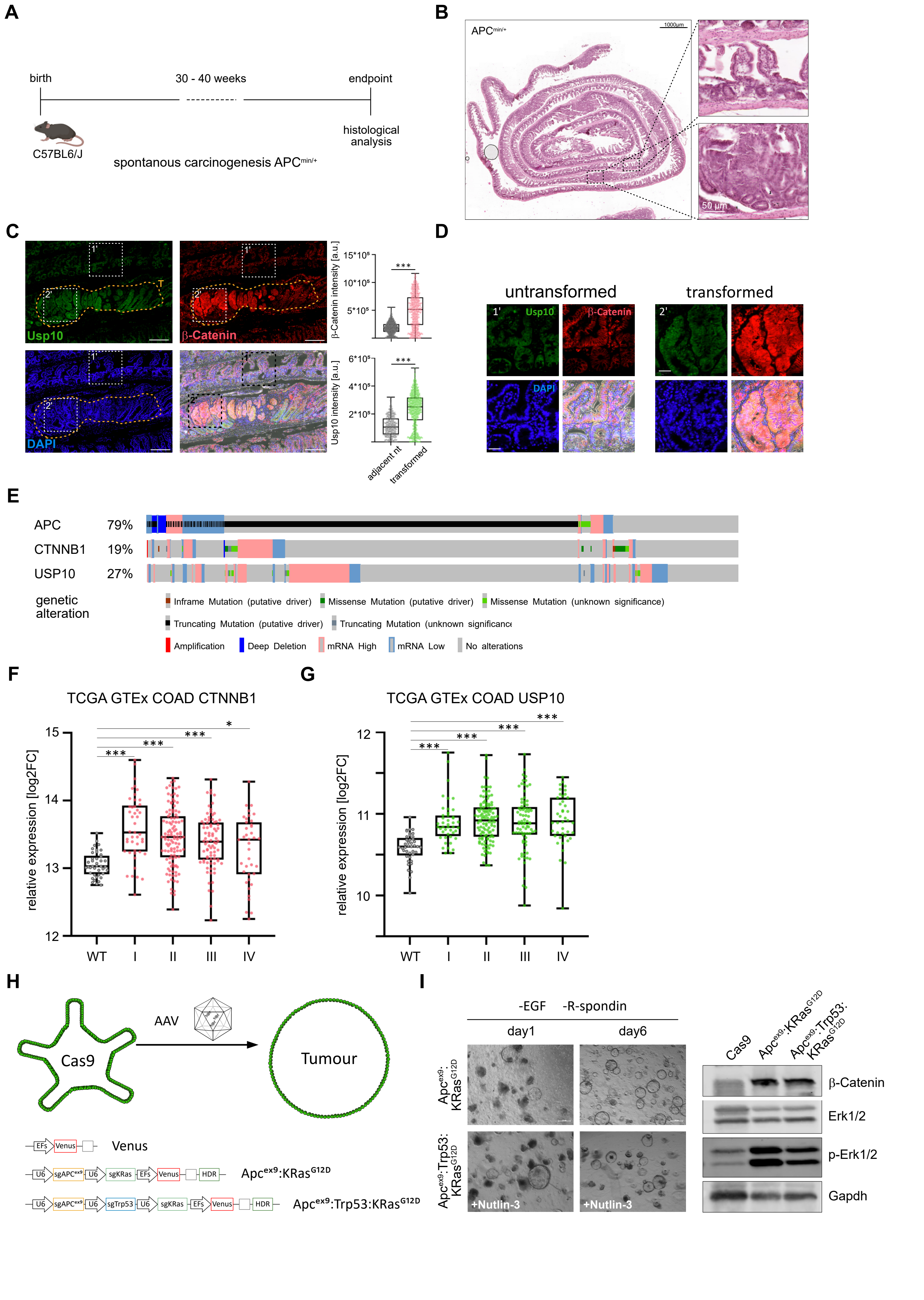


*Supplementary Figure 2*

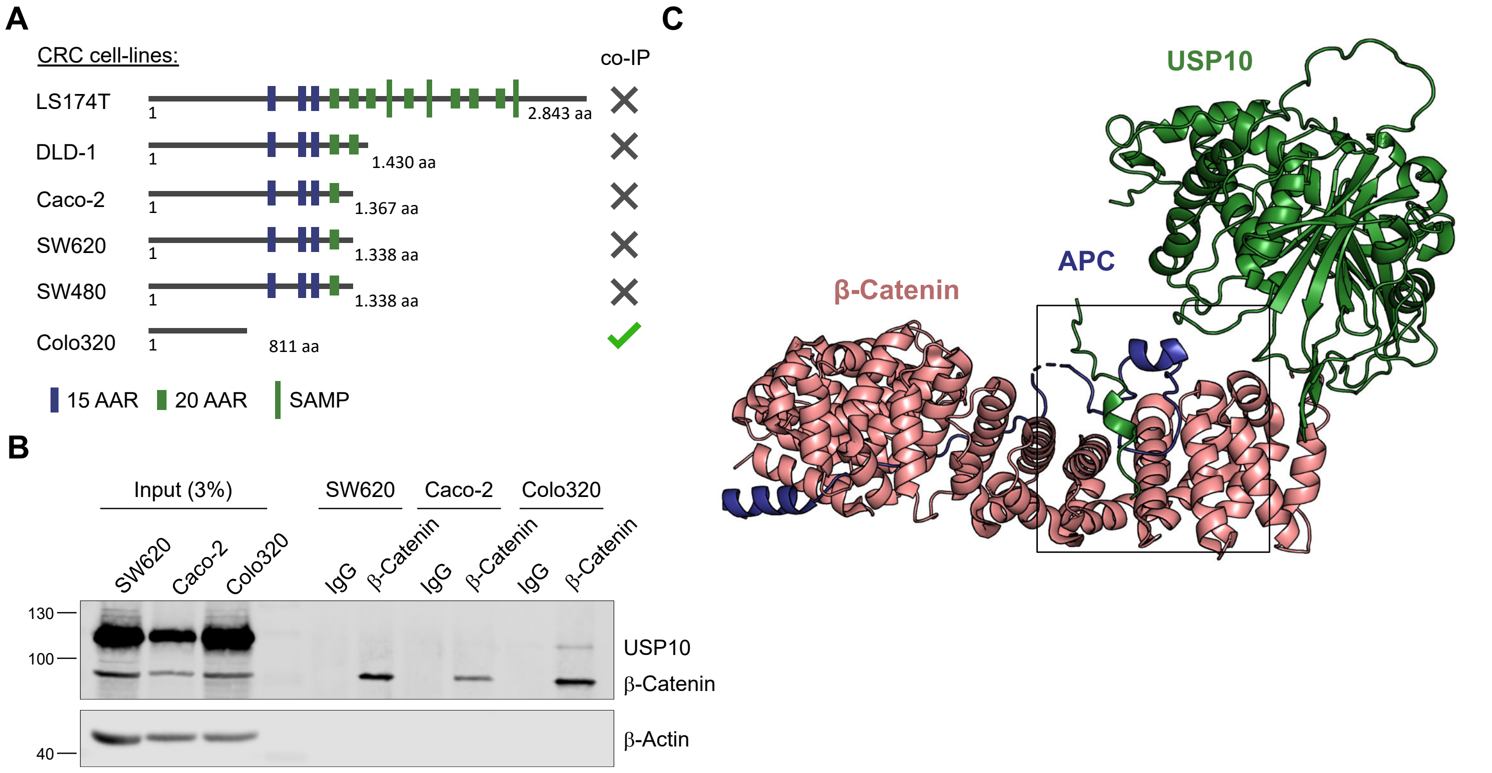


Supplementary Figure 3

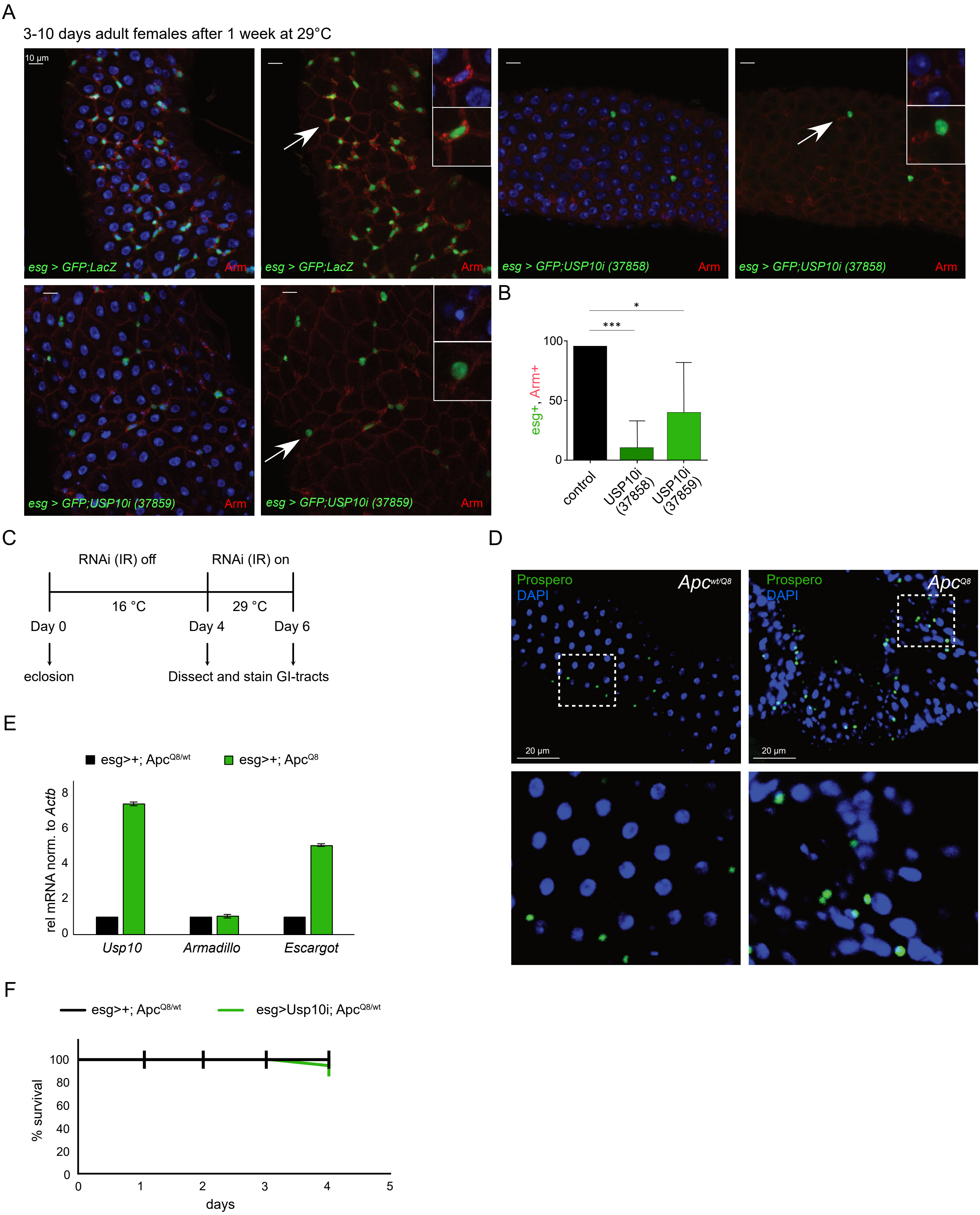


*Supplementary Figure 4*

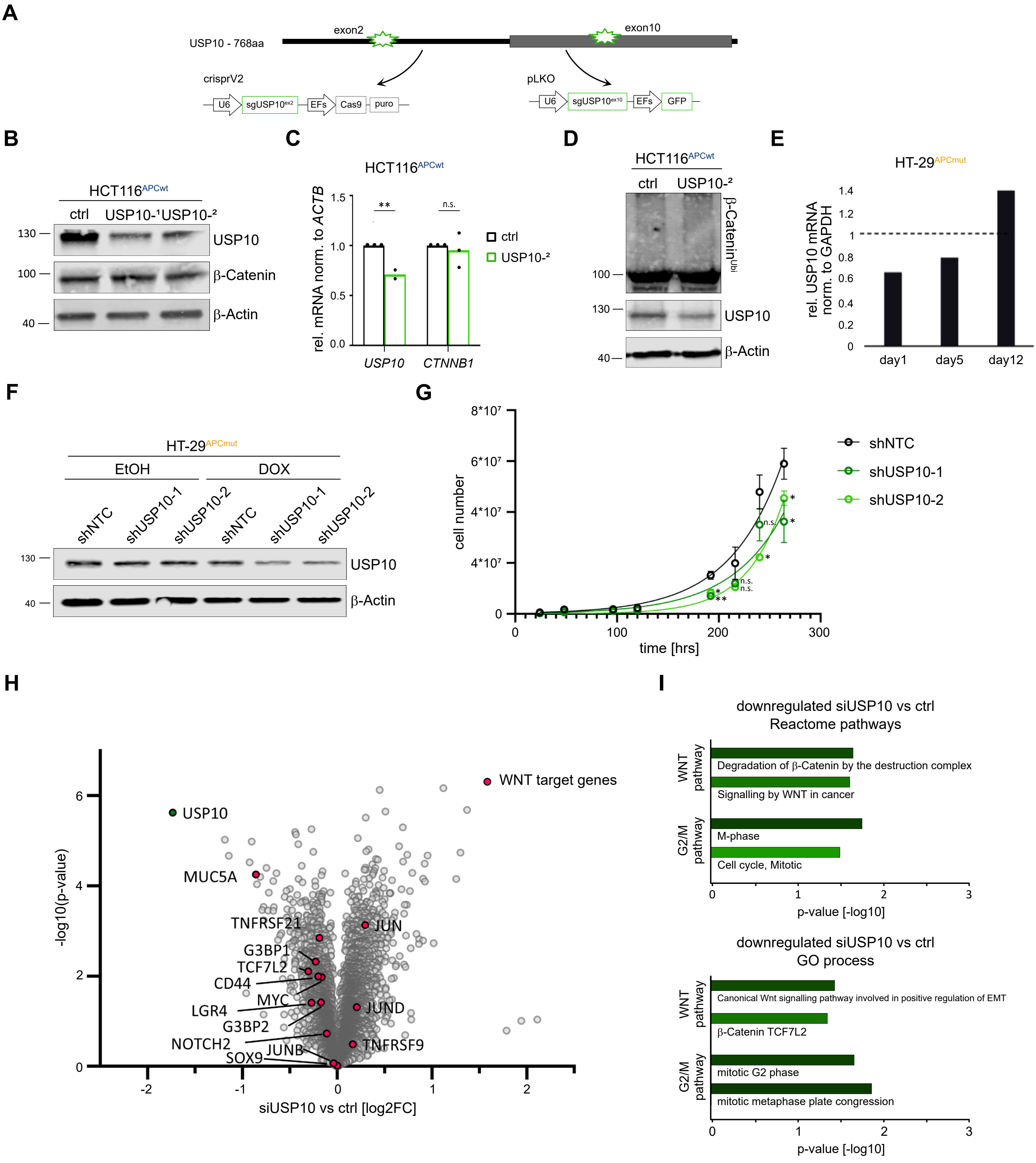


*Supplementary Figure 5*

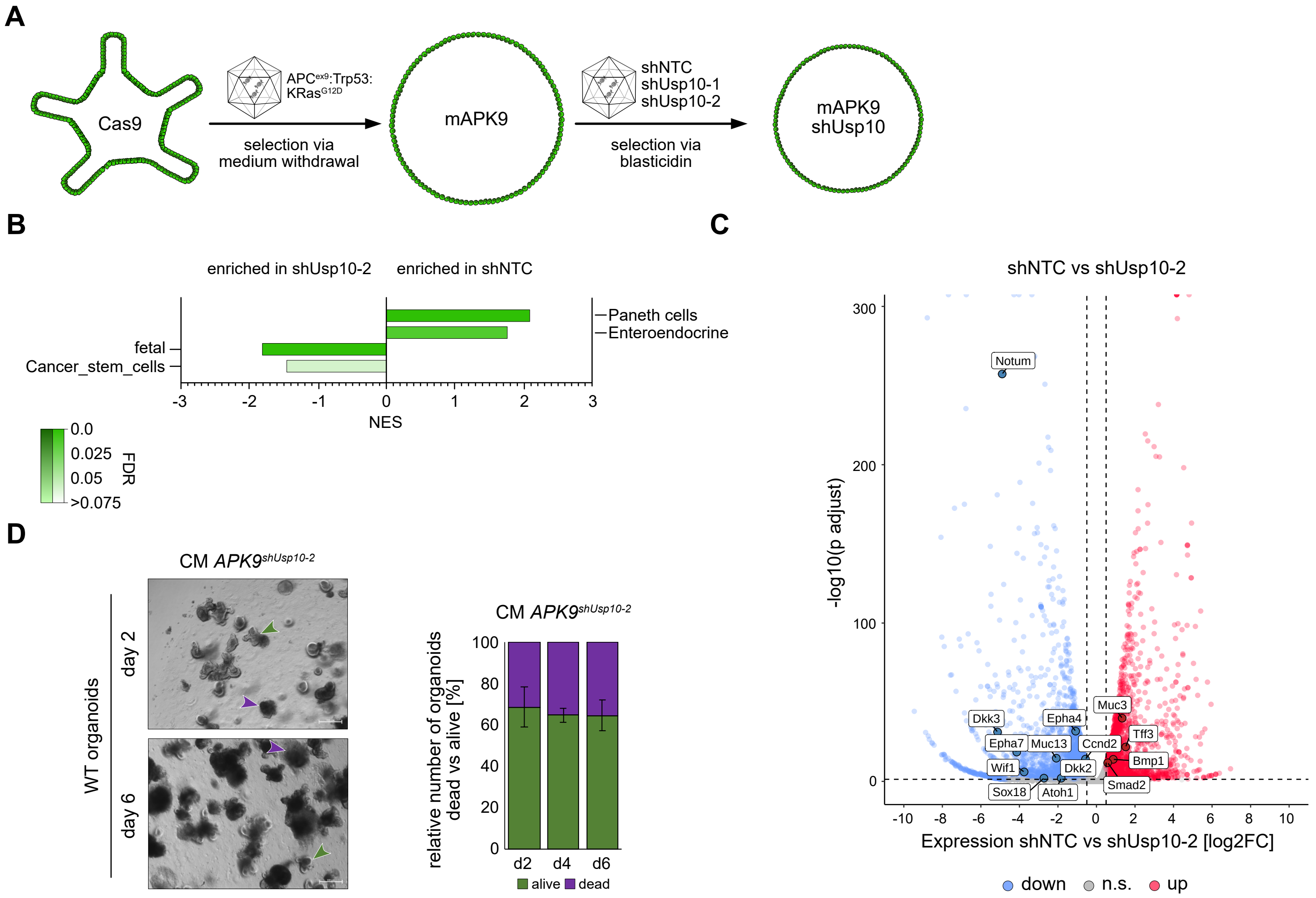


*Supplementary Figure 6*

**Supplementary Figure legends:**

Figure S1

A. Representative immunofluorescent images of endogenous β-Catenin (green) upon siRNA mediated knock-down of NTC (control) and the known negative regulators *USP8* and *USP15*. DAPI served as nuclear marker (blue).
B. Representative immunofluorescent images of endogenous β-Catenin (green) upon siRNA mediated knock-down of NTC (control) and the known positive regulators *UCHL1* and *USP4*. DAPI served as nuclear marker (blue).
C. Schematical model of a crypt with markers for Paneth cells (Lysozyme) and stem cells (Cd44).
D. Representative fluorescent images of WT intestinal crypt and villus. Left: endogenous Usp10 (green) and Lysozyme (magenta). Right: endogenous Usp10 (green) and Cd44 (magenta). DAPI served as nuclear marker (blue).

Figure S2

A. Schematic representation of the spontaneous CRC model utilising C57BL6/J $Apc^{min/wt}$ mice. Loss of heterozygosity and CRC onset in the small intestine and colon appears withing 30-40 weeks post birth.
B. Haematoxylin and eosin (H&E) staining of intestines of $Apc^{min/wt}$ mice 30 weeks post birth. Insets highlight either non-transformed adjacent tissue or primary tumour.
C. Representative immunofluorescent images and quantification of intestines of $Apc^{min/wt}$ mice 30 weeks post birth, of endogenous Usp10 (green) and β-Catenin (red), respectively. DAPI served as nuclear marker (blue). Insets highlight either untranformed (1') or transformed (2') regions. P-values were calculated using Mann-Whitney U test. ***p <0.001.
D. Insets from Figure S1G. High magnification immunofluorescent images of intestines of genetically engineered $APC^{min/+}$ mice for Usp10 (green) and β-Catenin (red). DAPI served as nuclear marker (blue).
E. Overview and percental occurrence of *APC, CTNNB1 and USP10* genetic alterations in CRC patients. Plot was generated using TCGA PanCancer data set using cBioportal.
F. +G. Expression of *CTNNB1* (A) *and USP10* (B) in non-transformed (WT) and stage I – IV CRC samples. Publicly available data were extracted from *UCSC Xena*: GTEx (n=41) and COAD (Stage I n = 44, II = 111, III = 81, IV = 41). P-values were calculated using Mann-Whitney test. *p<0.05; ***p<0.001
H. Schematic representation of Cas9 organoid infection strategy via AAV and expected morphological change upon organoid transformation. sgRNAs targeting Apc in exon 9, Trp53 and KRas with corresponding HDR-template under the control of the U6-Promoter.
I. Representative brightfield images of infected organoids 1- and 6- days post infection. Selection of successfully infected organoids was achieved via withdrawal of distinct medium-components. ENR - EGF, Noggin, R-spondin. For $Trp53^{\Delta}$ selection, 10 µM of the MDM2 inhibitor Nutlin-3 was used for six days. Immunoblot analysis of infected AK and APK organoids compared to control. β-Catenin and p-

ERK1/2 served as downstream targets for Apc truncation and KRas activation, respectively. Gapdh served as loading control.

Figure S3

A. Schematic representation of truncating mutations reported in the *APC* gene in the CRC cell lines LS174T, DLD-1, Caco-2, SW480, SW620 and Colo320 and summary about observed co-IP of USP10 and β-Catenin. Dark blue box = 15 AAR domains, green small boxes = 20 AAR domains, large green boxes = SAMP domains. 15- and 20-AAR = β-Catenin amino acid repeats; SAMP = Axin binding sites.
B. Representative input and endogenous co-immunoprecipitation of USP10 and β-Catenin in human CRC cell lines SW620, Caco-2 and Colo320 (APCmut). IgG served as antibody specificity control. β-Actin served as loading control. Input represents 3% of total loading. n=1.
C. Comparison of the AF2M predicted β-Catenin/complex and the crystal structure of β-Catenin bound to phosphorylated APC (PDB: 1TH1). USP10 (green), β-Catenin (salmon) and APC (blue). Highlighted is the common binding site on β-Catenin in which USP10 and APC are competing for binding.

Figure S4

A. Confocal images of adult females' midguts expressing UAS-GFP and the indicated transgenes under the control of the Escargot-Gal4 (Esg, GFP) promoter, that direct *GAL4* expression to progenitor cells. DAPI (blue) marks nuclei and arrows point to cells shown in insets. Elimination of USP10 in progenitor cells using the indicated UAS-RNAi lines, but not control, results in reduced number of progenitor cells and reduced progenitors expressing Armadillo (GFP+, Arm+, red).
B. Quantification of esg+ and Arm+ cells from A. Error bars represent standard deviation of n=3 independent experiments. Significance was calculated using students t-test. *p<0.05; ***p<0.001.
C. Schematic overview of the intestinal hyperplasia survival study using the heat shock induced siRNA expression in *D.melanogaster*.
D. Immunofluorescent images against endogenous Prospero (green) from midguts of $Apc^{Q8/+}$ or $Apc^{Q8/Q8}$ flies. Prospero was used to distinguish between big and polyploid EC-like cells, small Prospero- (ISC/EB) and small Prospero+ (EE) cells. DAPI served as nuclear marker (blue).
E. qRT-PCR analysis of the expression of d*Usp10*, *Armadillo* and *Escargot* in midguts isolated from either $Apc^{Q8/+}$ or $Apc^{Q8/Q8}$ flies. Rel. mRNA was normalised to *Actb*. Error bars represent standard deviation of n=3. Significance was calculated using unpaired t-test. *p-value<0.05; **p-value<0.005; ***p-value<0.001.
F. Kaplan Meier plot of adult *D.melanogaster* survival of the indicated genotypes. n=35.

Figure S5

A. Schematic model of *USP10* targeting strategy via CRISPR/Cas9. A double guide approach, targeting exon2 and 10 of *USP10*, with GFP and puromycin for selection was utilised.
B. Representative immunoblot against endogenous USP10 and β-Catenin in APC wild-type HCT116 cells upon CRISPR mediated depletion of USP10. Two different cell pools ($USP10^{-1}$, $USP10^{-2}$) along with non-targeting control cells are shown. β-Actin served as loading control. n=2.
C. Quantitative RT-PCR of *USP10* and *CTNNB1* expression of HCT116 $USP10^{-2}$. Error bars represent standard deviation of n=3 independent experiments. Significance was calculated using students t-test. **p<0.005; n.s.=non-significant.
D. Tandem Ubiquitin Binding Entity (TUBE) assay of endogenous poly-ubiquitylated proteins, followed by immunoblotting against endogenous β-Catenin in HCT116 $USP10^{-2}$. β-Actin served as loading control. n=2.
E. Quantitative RT-PCR of *USP10* expression in HT-29 $USP10^{\Delta}$ cells over time.
F. Immunoblot for two different DOX-inducible shRNAs against *USP10* and control shRNA in HT-29 cells after 4 days of 0.5 µM DOX and EtOH control treatment. Endogenous USP10 and β-Catenin was blotted. β-Actin served as loading control.
G. Growth-curve of shUSP10-1 and -2 compared to shNTC HT-29 cells. Error bars represent standard deviation of n=3 independent experiments. Significance was calculated using multiple paired t-tests. *p<0.05; **p<0.005; n.s.=non-significant.
H. Volcano-plot of up- and downregulated proteins upon siRNA mediated knock-down of USP10 in HT-29 cells. Proteins related to WNT signalling are highlighted in pink.
I. PANTHER pathway analysis from (K).

Figure S6

A. Schematic representation of two-step Cas9 organoid infection strategy via AAVs. sgRNAs targeting *Apc* in exon 9, *Trp53* and *KRas* with corresponding HDR-template under the control of the U6-Promoter were used to transform Cas9 organoids. Successfully selected APK organoids were then infected with AAV carrying shUsp10-1, -2 and shNTC and selected via blasticidin.
B. Gene set enrichment analysis of intestinal specific gene sets, deregulated in shUsp10-2 compared to shNTC APK9 organoids.
C. Volcano-plot of differential expressed genes in shUsp10-2 compared to shNTC organoids. Up- and down-regulated genes are highlighted in red and blue, respectively. Genes-of-interest are labelled.
D. Representative brightfield images of wild-type (WT) organoids cultured in $APK^{shUsp10-2}$ conditioned medium (CM), supplemented with EGF and R-spondin, for up to 6 days. Purple arrows indicate dead organoids, green arrows indicate live organoids. Quantification: Dead and alive organoids were counted and bar graphs represent percentage of alive vs dead organoids. Error bars represent standard deviation calculated from n=3 independent experiments.
