## Supplementary material for "Stabilisation of β-Catenin-WNT signalling by USP10 in APC-*truncated* colorectal cancer drives cancer stemness and enables super-competitor signalling": Reissland et al Material and Methods

### Consumables and Resources:

| 1st Antibodies | Company | Identifier |
|---|---|---|
| USP10 | Sigma-Aldrich | HPA006731 |
| B-ACTIN (C4) | Santa Cruz | sc-47778 |
| VINCULIN | Sigma-Aldrich | V9131 |
| c-MYC (N-262) | Santa Cruz | sc-764 |
| c-JUN (G-4) | Santa Cruz | sc-74543 |
| NOTCH1 | Bimake | A5176 |
| CTNNB1 | BD | 9315374 |
| LGR4 | Santa Cruz | sc-390630 |
| LGR5 | Thermo Fischer | MA5-25644 |
| OLFM4 | Thermo Fischer | PA5-73003 |
| ASCL2 | Thermo Fischer | PA5-40474 |
| CD44 | PTGLab | 60224-1-ig |
| KRT20 | Santa Cruz | sc-25725 |
| p-ERK (E-4) | Santa Cruz | sc-7383 |
| KI67 | Santa Cruz | sc-23900 |
| LAMIN A/C Alexa Fluor 790 | Santa Cruz | sc-376248 AF790 |
| TUBULIN | PTGLab | 60004-1-lg |
| ERK1/2 | Santa Cruz | sc-135900 |
| Delta | DHSB | C594.9B |
| Armadillo | DHSB | DCAD2 |

| 2nd Antibodies | Company | Identifier |
|---|---|---|
| SuperBoost™ Goat anti-Mouse Poly HRP | ThermoFisher | B40961 |
| SuperBoost™ Goat anti-Rabbit Poly HRP | ThermoFisher | B40962 |
| Donkey anti-Mouse IgG (H+L) Highly Cross-Adsorbed Secondary Antibody, Alexa Fluor 488 | ThermoFisher | A21202 |
| Donkey anti-Rabbit IgG (H+L) Highly Cross-Adsorbed Secondary Antibody, Alexa Fluor 488 | ThermoFisher | A21206 |
| Donkey anti-Mouse IgG (H+L) Cross-Adsorbed Secondary Antibody, DyLight 680 | ThermoFisher | SA5-10170 |
| Donkey anti-Goat IgG (H+L) Cross-Adsorbed Secondary Antibody, DyLight 680 | ThermoFisher | SA5-10090 |
| Donkey anti-Rabbit IgG (H+L) Cross-Adsorbed Secondary Antibody, DyLight 800 | ThermoFisher | SA5-10044 |
| Donkey anti-Mouse IgG (H+L) Cross-Adsorbed Secondary Antibody, DyLight 800 | ThermoFisher | SA5-10172 |
| Donkey anti-Goat IgG (H+L) Cross-Adsorbed Secondary Antibody, DyLight 800 | ThermoFisher | SA5-10092 |

| **Chemicals and Commercial Assays** | **Company** | **Identifier** |
|---|---|---|
| Gibco™ Dulbecco's Modified Eagle Medium (DMEM), high glucose | ThermoFisher | 11574486 |
| Gibco™ LHC Basal Medium (1X) | ThermoFisher | 11524556 |
| Gibco™ Trypsin-EDTA (0.5%), No Phenol Red | ThermoFisher | 15400054 |
| Fetal Bovine Serum (FCS) | Sigma-Aldrich | 12103C |
| Penicillin-Streptomycin | Sigma-Aldrich | P4333 |
| Polybrene | Sigma-Aldrich | TR-1003 |
| Polyethylenimine, Linear, MW 25000, Transfection Grade (PEI 25K) | Polysciences | 23966-1 |
| Dimethyl sulfoxide (DMSO) | Sigma-Aldrich | D8418 |
| Ethanol (Etoh) | Carl Roth | 5054.6 |
| Gibco™ Phosphate-buffered saline (PBS) | ThermoFisher | 10010031 |
| Nuclease-free water | Merck | 3098 |
| Pierce™ Protein A/G Magnetic Beads | ThermoFisher | 88802 |
| 2-Propanol/ Isopropanol | ROTH | AE73.2 |
| Adenosintriphosphat (ATP) | Jena Bioscience | NU-1010-10G |
| Agarose | ROTH | 3810.4 |
| Ampicillin (Amp) | ROTH | HP62.2 |
| Bovine serum albumine (BSA) | Merck Millipore | 810683 |
| Dithiothreitol (DTT) | Sigma-Aldrich | D9779 |
| Eosin | Sigma | E4009 |
| Hematoxylin | Sigma | H3136 |
| Polyvinylidene difluoride membranes (PVDF) Immobilon Transfer Membrane | Merck | IPFL00010 |
| N,N,N',N'-tetramethylenethylendiamine (TEMED) | ROTH | 2367.3 |
| Natrium chloride (NaCl) | AppliChem | A2942,1000 |
| Neutrally buffered formalin (NBF) | Thermo Fisher | 5700TS |
| Tris-HCl | ROTH | 9090.5 |
| TritonX100 | ROTH | 3051.3 |
| Xylene | Sigma | 534056 |
| β-Mercaptoethanol | ROTH | 4227.1 |
| Methanol (MeOH) | ROTH | 0082.3 |
| Random Hexamer Primer | ThermoFisher | SO142 |
| RiboLock RNase Inhibitor | ThermoFisher | EO0381 |
| Histo-Clear® Histological Clearing Agent | National Diagnostics | HS-200; Lot03-19-18 |
| Cytoseal 60™ | Thermo Scientific | 8310-4 |
| SignalStainR DAB Substrate Kit | Cell Signaling | 8059 S |
| Doxycycline hyclate | Sigma-Aldrich | D9891 |
| Propidium Iodide (PI) | Sigma-Aldrich | 11348639001 |
| 4′,6-diamidino-2-phenylindole (DAPI) | ThermoFisher | D1306 |
| Hoechst | ThermoFisher | 62249 |

| Experimental Models: Organisms/Strains Cell | Company | Identifier |
|---|---|---|
| B6(C)-Gt(ROSA)26Sor$^{em1.1(CAG-cas9*,-EGFP)Rsky}$/J | The Jackson laboratory | Stock No: 028555 |
| C57BL/6J | The Jackson Laboratory | Stock No: 000664 |
| Balb/c | Charles River | Stock No: 028 |
| **Oligonucleotides and recombinant DNA** | **Sequence** | **Company** |
| sgRNA_USP10_ex2 fwd | caccgCTTACCTCAACTGAAG | Sigma |
| sgRNA_USP10_ex10 fwd | caccGCTCTTCTTCATTGACCGAG | Sigma |
| shRNA hUSP10 #1 | caccgaccagcaacaacacttgtaaaTTCAAGAGAtttacaagtgttgttgctggtcTTTTT | Sigma |
| shRNA hUSP10 #2 | caccgcctatgtggaaactaagtattTTCAAGAGAaatacttagtttccacataggcTTTTT | Sigma |
| shRNA mUsp10 # 1 | caccgtcattagagatcgctcttacTTCAAGAGAgtaagagcgatctctaatgacTTTTT | Sigma |
| shRNA mUsp10 # 2 | caccgcacagcctacctcctatattTTCAAGAGAaatataggaggtaggctgtgcTTTTT | Sigma |
| sgRNA mKras | GACTGAGTATAAACTTGTG | Sigma |
| sgRNA mApc Exon9 | caccGCCGCTAGAACTCAAAACAC | Sigma |
| sgRNA mTrp53 | CACCGATGGTGGTATACTCAGAGC | Sigma |

| | | |
|---|---|---|
| mKrasG12D repair template | TCCATGTATTTTTATTAAGT<br>GTTGATGAGAAAGTTGTAA<br>GTGACTTACAGGTTACTCT<br>GTACATCTGTAGTCACTGA<br>ATT<br>CGGAATATCTTAGAGTTTT<br>ACACACAAAGGTGAGTGTT<br>AAAATATTGATAAAGTTTT<br>TGATAATCTTGTGTGAGAC<br>ATGT<br>TCTAATTTAGTTGTATTTTA<br>TTATTTTTATTGTAAGGCCT<br>GCTGAAAATGACTGAGTAT<br>AAACTTGTGGTaGTTGGAG<br>CT<br>GatGGCGTAGGCAAGAGCG<br>CCTTGACGATACAGCTAAT<br>TCAGAATCACTTTGTGGAT<br>GAGTATGACCCTACGATAG<br>AGGT<br>AACGCTGCTCTACAGTCTG<br>CGTGCGCTTGTAAAGGACG<br>GCAGCCAGCCGCTTTGAAA<br>AAGATATCATTTTTATATTT<br>ATT<br>AGAAAATTATATTGAAAGT<br>TATTTCAGTTATATGTGAT<br>GTCCTTTAGTTCCAAGGCT<br>TTAAACTGGGTGT | IDT |
| pLKO5.sgRNA.EFS.GFP | Addgene 57822 | Addgene plasmid # 57822 |
| pLKO shUSP_10_1 (GFP) | [21] | N/A |
| pLKO shUSP_10_2 (GFP) | [21] | N/A |
| pHelper | Cell Biolabs, INC. | VPK-400-DJ |
| pAAV-DJ Vector | Cell Biolabs, INC. | VPK-420-DJ |
| AAV:ITR- U6-sgRNA(p53)- pEFS-2A-mCherry-shortPA- ITR | [21] | N/A |
| pHelper | Cell Biolabs, INC. | VPK-400-DJ |
| pAAV-DJ Vector | Cell Biolabs, INC. | VPK-420-DJ |
| pUMVC | pUMVC was a gift from Bob Weinberg | Addgene plasmid # 8449 |
| pCMV-VSV-G | pCMV-VSV-G was a gift from Bob Weinberg | Addgene plasmid # 8454 |

| | |
|---|---|
| Fly lines | Q462 esg-GAL4 UAS-GFP/CyO; tub-GAL80[ts]/TM6B<br>Q469 esg-GAL4 UAS-GFP/CyO; Apc1[Q8] FRT82B/TM6B<br>37859-vdrc w; UAS-Usp10-IR [GD]/TM3<br>y w; UAS-Usp10-IR/CyO; FRT Apc1[Q8]/TM6B<br>Bloomington-50800 y w; Usp10:GFP/TM3, Ser Sb<br>vdrc-37858 w; UAS-Usp10-IR [GD; 2nd] |

| Software | Company/Source |
|---|---|
| cBioportal | https://www.cbioportal.org |
| GEPIA and GEPIA2 | http://gepia.cancer-pku.cn |
| KM-plotter | http://kmplot.com/analysis/ |
| Operetta Imaging | Perkin Elmer |
| BoxPlotR | http://shiny.chemgrid.org/boxplotr/ |
| Morpheus | https://software.broadinstitute.org/morpheus/ |
| Excel | Microsoft |
| Affinity Desgner | h https://affinity.serif.com/es/designer/ |
| Image Studio | Licor |
| Panther Classification system | http://pantherdb.org |
| AATBIO IC50 calculator | https://www.aatbio.com/tools/ic50-calculator |
| PRISM4 | GraphPad Software, Inc. |
| Affinity Designer | Serif Europe |
| ImageJ | National Insistute of Health |
| Primerx | http://www.bioinformatics.org/primerx/cgi-bin/DNA_1.cgi |
| ROC Plotter | http://www.rocplot.org/ |
| Pannoramic Case Viewer | 3dHistech |
| R2: Genomics Analysis and Visualization Platform | http://r2.amc.nl |
| UCSC Xena | https://ucsc-xena.gitbook.io/project/ |
| Proteome discoverer 2.2 | Thermo Scientific |

| | |
|---|---|
| MaxQuant | https://www.maxquant.org/ |
| Perseus 1.6.5. | https://maxquant.net/perseus/ |
| Uniprot | https://www.uniprot.org/ |
| INSTANT CLUE | https://kups.ub.uni-koeln.de/17610/ |
| FastQC | http://www.bioinformatics.babraham.ac.uk/projects/fastqc/ |
| Bowtie2 v2.3.4.1 | http://bowtie-bio.sourceforge.net/index.shtml |
| TopHat v.2.1.1 | https://ccb.jhu.edu/software/tophat/index.shtml |
| Samtools v1.3 | http://samtools.sourceforge.net |
| R | https://www.r-project.org |
| EdgeR | https://bioconductor.org/packages/release/bioc/html/edgeR.html |
| COMBENEFIT | https://www.cruk.cam.ac.uk/research-groups/jodrell-group/combenefit |
| Harmony Software | Perkin Elmer |
| EMBL | https://www.embl.de/ |
| SPLASHRNA | http://splashrna.mskcc.org/ |
| BD FACSDiva 6.1.2 BD | Biosciences |
| FlowJo 8.8.6 | FlowJo, LLC |
| Illustrator TM, | Adobe Inc. |
| Photoshop TM, | Adobe Inc. |
| Acrobat TM | Adobe Inc. |
| Integrated Genome Browser | Nicol et al. 2009 |
| Mac OS X | Apple Inc. |
| Office 2011 Mac | Microsoft Inc. |
| Qupath | https://qupath.github.io |
| UCSC Genome Bioinformatics | http://genome.ucsc.edu |
| ApE plasmid editor | By Wayne Davis |
| Venn diagrams | http://bioinformatics.psb.ugent.be/webtools/Venn/ |
| Nemates | http://nemates.org |
| Online Web statistical calculator Astatsa | https://astatsa.com/ |

| | |
|---|---|
| DOI citation formatter | https://citation.crosscite.org |
| Zhang lab gRNAs design resources | https://zlab.bio/guide-design-resources |
| CHOPCHOP | http://chopchop.cbu.uib.no/ |
| RNAi Consortium | www.broadinstitute.org/rnai-consortium/rnai-consortium-shrna-library |
| Cancer Therapeutics Response Portal | https://portals.broadinstitute.org/ctrp.v2.1/ |
| Genomics of Drug Sensitivity in Cancer | https://www.cancerrxgene.org/ |
| Catalogue Of Somatic Mutations In Cancer (COSMIC) | https://cancer.sanger.ac.uk/cell_lines |
| Gene Expression and Mutations in Cancer Cell Lines (GEMiCCL) | https://www.kobic.kr/GEMICCL/ |
| Clust Vis | https://biit.cs.ut.ee/clustvis/ |
| Depmap | https://depmap.org/portal/ |
| **Instrument** | **Company** |
| Odyssey® CLx Imaging System | Licor |
| iBright™ FL1000 Imaging System | Invitrogen |
| BD FACSCanto II Cell Analyzer | BD Biosciences |
| StepOnePlus Real-Time PCR System | Thermo Scientific |
| Invitrogen Countess II FL Automated Cell Counter | Thermo Scientific |
| Pannoramic DESK scanner | 3DHISTECH |
| FSX100 microscopy | Olympus Life Science |
| Operetta High-Content Imaging System | Perkin Elmer |

| | |
|---|---|
| Fragment Analyzer | Agilent formerly Advanced Analytical |
| Axiocam 503 mono | Zeiss |
| Branson Sonifier 250 | Branson |
| 250 mm long C18 column: X-Bridge, 4.6 mm ID, 3.5 µm particle size | Waters |
| EASY-nLC™ 1200 System | Thermo Scientific |
| Orbitrap Fusion Lumos mass spectrometer | Thermo Scientific |
| 1.9 µm C18 particles | ReproSil-Pur, Dr. Maisch |
| Hyrax M55 Rotary Microtome | Leica |
| Mr. Frosty freezer container | Thermo Scientific |
| PCR cycler: SimpliAmp thermo cycler | Life technologies |
| Experion™ Automated Electrophoresis System | Bio-Rad |
| Cell culture incubator BBD 6220 | Heraeus |
| Casy® cell counter | Innovatis |
| Centrifuge Avanti J-26 XP | Backman Coulter |
| Centrifuge Eppendorf 5417 R | Eppendorf |
| Centrifuge Eppendorf 5425 | Eppendorf |
| Centrifuge Eppendorf 5430 | Eppendorf |
| Centrifuge Galaxy MiniStar | VWR |

| | |
|---|---|
| Centrifuge Multifuge 1S-R | Heraeus |
| Deep-sequencer Genome Analyzer IIx | Illumina |
| Dry Bath System | Starlab |
| Thermomixer® comfort | Eppendorf |
| Incubator shaker Model G25 | New Brunswick Scientific |
| Luminometer GloMax | Promega |
| Microscopes Axiovert 40CFL | Zeiss |
| PCR thermal cycler Mastercycler pro S | Eppendorf |
| Spectrofluorometer NanoDrop 1000 | Thermo Scientific |
| UltrospecTM 3100 pro UV/Visible | Amersham Biosciences |
| SDS page system Minigel | Bio-Rad |
| SDS page system Tetra Cell | Bio-Rad |
| Maxi UV fluorescent table | Peqlab |
| Mixer Vortex-Genie 2 | Scientific Industries |
| Julabo ED-5M water bath | Julabo |
| Memmert waterbath | Memmert |
| Immunoblot transfer system: Perfect Blue Tank Electro Blotter Web S | Peqlab |
| Power supply: Power Pac | Bio-Rad |

| | |
|---|---|
| Chemiluminescence imaging LAS-4000 mini Fujifim | Fujifim |
| Sterile bench HeraSafe | Heraeus |
| Siemens linear accelerator for X-ray irradiation | Siemens |
| Pipetman Classic P2.5, P10, P20, P200 and P1000 | Gilson |
| Dionex Ultimate 3000 analytical HPLC | Thermo Scientific |
| Leica VT 1200S | Leica |
| Microscope TCS SP5 | Leica |
| BD FACS Aria III | BD Biosciences |
| Pipetboy acu 2 | Integra |
| Consort EV243 electrophoresis power supply | Sigma |
| Ventana DP 200 slide scanner | Roche |

| **Cell lines** | **Source** | **Identifier** |
|---|---|---|
| Human: HEK 293T | ATCC | CRL-11268 |
| HT-29 | ATCC | HTB-38 |
| Colo 320 | ATCC | CCL-220 |
| DLD-1 | ATCC | CCL-221 |
| SW 480 | ATCC | CCL-228 |
| SW 620 | ATCC | CCL-227 |
| HCT 116 | ATCC | CCL-247 |
| Ls174T | ATCC | CL-188 |
| **RT-PCR primers** | **Sequence** | **Company** |
| hLGR5_fw | CTCCCAGGTCTGGTGTGTTG | Sigma |
| hLGR5_rev | GAGGTCTAGGTAGGAGGTGAAG | Sigma |
| hOLFM4_fw | TTCTCCTAGCCCTTCTGTTCTTCC | Sigma |

| | | |
|---|---|---|
| hOLFM4_rev | TTCCAAGCGTTCCACTCTGTCC | Sigma |
| hDCLK1_fw | GGTGAACGTCAAGA CCACCT | Sigma |
| hDCLK1_rev | GTCCTGAAGGCACATCACCT | Sigma |
| hALDH1_fw | TGTTAGCTGATGCCGACTTG | Sigma |
| hALDH1_rev | TTCTTAGCCCGCTCAACACT | Sigma |
| hCCND1_fw | CTCCGCCTCTGGCATTTTGG | Sigma |
| hCCND1_rev | TCTCCTTGCAGCTGCTTAG | Sigma |
| hCD44_fw | AGAAGGTGTGGGCAGAAGAA | Sigma |
| hCD44_rev | AAATGCACCATTTCCTGAGA | Sigma |
| hc-JUN_1_fw | TTCTATGACGATGCCCTCAACGC | Sigma |
| hc-JUN_1_rev | GCTCTGTTTCAGGATCTTGGGGTTAC | Sigma |
| hc-JUN_2_fw | CAAGACCTTCCTCTCAGGCTCATTC | Sigma |
| hc-JUN_2_rev | GGGGCATGTTTGGAGGGACA | Sigma |
| mOLFM4_fw | GCCACTTTCCAATTTCAC | Sigma |
| mOLFM4_rev | GAGCCTCTTCTCATACAC | Sigma |
| mLGR5_fw | CGAGCCTTACAGAGCCTGATACC | Sigma |
| mLGR5_rev | TTGCCGTCGTCTTTATTCCATTGG | Sigma |
| mSOX9_fw | CTGGAGGCTGCTGAACGAGAG | Sigma |
| mSOX9_rev | CGGCGGACCCTGAGATTGC | Sigma |
| mASCL2_fw | CCTCTCTCGGACCCTCTCTCAG | Sigma |
| mASCL2_rev | CAGTCAAGGTGTGCTTCCATGC | Sigma |
| mMuc2_fw | TGGAGCCTGAAACACAATCACT | Sigma |
| mMuc2_rev | ATGAAAATCAACAACCAGCTCATCT | Sigma |
| mCD44_fw | GTTGATGGCTCCTTACCAGG | Sigma |
| mCD44_rev | GGTTCGCACTTGAGTGTCCAG | Sigma |
| mc-Jun_fw | GCGGCTGCAAGCCCTGAAG | Sigma |
| mc-Jun_rev | GAGACTCCATGTCGATAG | Sigma |
| mAxin2_fw | CCACCAAGACCTACATAC | Sigma |
| mAxin2_rev | CCACTCCTCACATATTCC | Sigma |
| mCCND1_fw | CAGAGGCGGATGAGAACAA | Sigma |
| mCCND1_rev | AGGGTGGGTTGGAAATGAA | Sigma |
| mCTNNB1_fw | ACCAGAGTGAAAAGAACGGTAG | Sigma |
| mCTNNB1_rev | AATGGCTTGGAATGAGACTG | Sigma |

### Supplementary Material and Methods

#### Tissue culture, transfection, infection and reagents.

Cells were plated on Greiner petri dishes and maintained at 37°C, 95% relative humidity and 5% $CO_2$ for optimal growth conditions. All cell lines were obtained from ATCC or ECACC. HEK-293T, HCT116, HT-29, SW480 and SW620 cells were cultured in DMEM (Gibco) supplemented with 10% fetal bovine serum (FCS). LS174T, DLD-1, Caco-2 and Colo320 cells were cultured in RPMI 1640 (Gibco) supplemented with 10% FCS.

For DNA transfection, a mix of 2.5 µg plasmid DNA, 200 µl medium and 5 µl PEI was added into the 6-well dish medium (60 % confluence), after 6 hrs of incubation at 37°C the medium was changed to full supplemented medium.
For DNA infection, AAVs or Lentiviruses were added to the cell medium in the presence of polybrene (5 µg/ml) and incubating at 37°C for 96h. After incubation, infected cells were selected with 1 - 2 µg/ml Puromycin for 72h, 20 µg/ml Blasticidin for 1 weeks or FACS-sorted for GFP positive cells (FACS Canto II BD).

#### Murine tumour models

All *in vivo* experiments were approved by the Regierung Unterfranken and the ethics committee under the license numbers 2532-2-555, 2532-2-556, 2532-2-694 and 2532-2-1002. The mouse strains used for this publication are listed. All animals are housed in standard cages in pathogen-free facilities on a 12-h light/dark cycle with *ad libitum* access to food and water. FELASA2014 guidelines were followed for animal maintenance.

*Acute oncogenesis upon CRISPR mediated targeting of Apc in vivo*

Detachment of the intestinal mucus:
By administering 2%v/v Dextran Sulfate Sodium (DSS), dissolved in drinking water for three days prior to treatment detached the intestinal mucus layer sufficiently and

enhanced the accessibility of the stem cell niche. Post infection, animals were switched to regular drinking water and access given *ad libidum*.

Enema:
Mice were anaesthetised with isoflurane and an enema (500ul PBS) is administered by colorectal injection to flush the rectum. This also loosens newly formed mucus and makes the intestinal mucosa more accessible to viral infection. This method is also used in a similar form for patients. The animals are allowed to wake up from the anaesthesia again and the clistier is flattened over a period of 30 minutes.

Viral infection by enema:
This is followed by re-anaesthesia with isoflurane and lentiviral colorectal infection with a round-headed needle. The round-headed needle is inserted via the anus and the virus, dissolved in physiological PBS (p.H. 7.4), is applied with a volume of 500 µl per animal. The animals wake up from anaesthesia shortly after the treatment and can be moved to their cages.

Anaesthesia and analgesia:
Anaesthesia by inhalation of isoflurane. A concentration of 3% is administered for induction, then anaesthesia is maintained at 2% until the end of the experiment. The total duration of isoflurane anaesthesia is approximately 5 minutes per animal.

Animals were sacrificed by cervical dislocation 12 weeks post rectal infection and the intestinal lining was fixed using 5% NBF, followed by Swiss role preparation. For IHC, IF and H&E, slides were de-paraffinized and rehydrated following the protocol: 2x 5 min. Xylene, 2x 3 min. EtOH (100%), 2x 3 min. EtOH (95%), 2x 3 min. EtOH (70%), 3 min. EtOH (50%) and 3 min. $H_2O$. For all staining variants, slides were mounted with 200 µl of Mowiol® 40-88 covered up by a glass coverslip. IHC slides were recorded using Pannoramic DESK scanner and analysed using Case Viewer software (3DHISTECH), QuPath and ImageJ. IF samples were recorded using FSX100 microscopy system (Olympus)

*AOM/DSS*

Azoxymethane was dissolved in 0.9% NaCl (0.5mg/ml) and injected intraperitoneally (i.p.; 10 mg/kg) once weekly for six times. DSS (MP Biomedicals, 216011090) was dissolved in drinking water. 0.5 g/l L-NNA (Sigma, N5501) was dissolved in acidified drinking water (pH 6). All mouse experiments were reviewed and approved by the Regierungspräsidium Darmstadt, Darmstadt, Germany.

**Histopathology and human CRC TMA**

For IHC and H&E, slides were de-paraffinized and rehydrated as following: IHC slides were subjected to epitope retrieval and blocked in 3% BSA at RT for 1 h. The antibody manufacturer's instructions were followed for all antibodies. In general, primary antibodies (diluted in 1% BSA) were incubated overnight at 4 °C, followed by three washes with PBS and subsequent incubation with the DAB secondary antibody for 1 h at RT. Then, slides were washed twice with 1xPBS for 5 min and stained with the DAB staining solution in 1xPBS. Upon DAB staining, slides were counterstained with hematoxylin. Slides were mounted using 200 µl of Mowiol® 40–88 covered by a glass coverslip. IHC slides were recorded using a Panoramic DESK scanner or FSX100 microscopy system (Olympus) and analyzed using Case Viewer software (3DHISTECH), QuPath software, PRISM and ImageJ. Antibodies used are listed in the consumables section.

**Patient-derived tissue microarrays (TMA)**

Paraffin molds were cast using an Arraymold Kit (IHC World, Kit D, IW-115, core diameter 2 mm, 36 cores). Human samples were cut and stained using hematoxylin and eosin and digitalized using a 3D Histech slide scanner (panoramic FLASH). Tumor and non-transformed tissue were identified and manually 'punched' and transferred from the tissue block to the tissue array. Upon completion, 3-µm thick sections were cut using a microtome and processed as described above.

### Fly models

Experiments using adults females midguts were performed similar to previous reports[1]. In brief, fly stocks were maintained on yeast-cornmeal-molasses-malt extract medium at 18°C or as stated in the text. Conditional expression of transgenic lines in ISCs was achieved by activating a UAS-transgene under the expression of the progenitor-specific Gal4 line escargot:Gal4 driver together with the tub-Gal80ts construct[2]. Flies were raised at 18°C. 2-4 days old, F1 adult progeny were transferred to the restrictive temperature 29°C (Gal80 off, Gal4 on) for the indicated time, dissected and analyzed. At least three biological independent repeats were performed for each experiment. Gut dissection and immunofluorescence detection: Gut fixation and staining were carried out as previously described[3]. Antibodies: mouse anti-Delta (1:50) and anti-Armadillo (1:50) mAbs were from the developmental hybridoma bank.

### Intestinal Organoids

*Isolation of murine intestinal organoids*

Murine intestinal organoids were isolated and cultured as described by Sato et al. in 2009. In brief, mice were sacrificed and an approximately 4 to 5 cm long piece of the small intestine cut out. The piece of intestine was directly transferred to ice cold PBS in a petri-dish, washed and cut into small pieces. Crypt isolation was achieved using 25 mM EDTA (30 min, 4°C), pieces were washed and isolated crypts were plated in Geltrex™ droplets and covered with murine ENR (ADF base +++, N2, B27, EGF, Noggin, R-spondin) medium.

*Cultivation of organoids*

For passaging, organoids were re-supended in PBS, mechanically disrupted into small fragments by pipetting, washed and re-suspended in fresh geltrex and plated at the desired ratio. For freezing, the organoid-pellet was re-suspended in freezing medium (50 % medium, 40 % FCS and 10 % DMSO) and stored at −80 °C. For thawing, organoids were quickly thawed and transferred to a falcon with medium. To remove the DMSO from freezing medium, organoids were centrifuged for 5 min at 900 x g, the supernatant was discarded, and the remaining pellet was re-suspended in Geltrex and plated.

*Infection of organoids*

For AAV- infection of organoids the spin-occulation method was used. Organoids were washed twice with PBS, re-suspended in 450 µL medium, mixed with 50 µL concentrated virus and transferred to a 24-well ultra-low attachment plate. For spin-occulation the plate was centrifuged for 90 min at 900 x g and 37 °C, followed by 4 hrs incubation at 37°C. After incubation, the organoid-virus suspension was washed (5 min, 900 g), re-supended in Geltrex™ and plated. After two days, selection of infected organoids was started by adding corresponding antibiotics to the medium or withdrawal of growth factors.

**RT-PCR**

RNA was isolated with ReliaPrep™RNA Miniprep Systems, according to manufacturer`s instructions. RNA was reverse transcribed into complementary DNA (cDNA) using random hexanucleotide primers and M-MLV reverse transcriptase (Promega). Quantitative RT-PCR was performed using qPCR SYBR Green mix (Applied Biosystem) on the instrument Step One™ Plus cycler. (Applied Biosystem) RT-PCR was performed using the next Program: 95°C for 15 min., 40x [95°C for 15 sec., 60°C for 20 sec. and 72°C for 15 sec.], 95°C for 15 sec. and 60°C for 60 sec. Relative expression was generally calculated with ΔΔCt relative quantification method using the expression of a House-keeping gene. For all the experiments, melting curves were performed.

**Immunological Methods**

Cells were lysed in RIPA lysis buffer (20 mM Tris-HCl pH 7.5, 150 mM NaCl, 1mM $Na_2$EDTA, 1mM EGTA, 1% NP-40 and 1% sodium deoxycholate), containing proteinase inhibitor (1/1000) by sonication using Branson Sonifier 250 with a duty cycle at 20%, output control set on level 2 and the timer set to 1 min (10 sonication cycles per sample). Protein concentration was quantified using Bradford assay. After mixing 1 ml of Bradford reagent with 1µl of sample, the photometer was used to normalize the protein amounts with a previously performed bovine serum albumin (BSA) standard curve. The quantified protein (25 µg) was boiled in 5x Laemmli buffer (312.5mM Tris-HCl pH 6.8, 500 mM DTT , 0.0001% Bromphenol blue, 10% SDS and 50% Glycerol) for 5 min and separated on 10% Tris-gels in Running buffer (1.25M Tris base, 1.25M glycine and 1% SDS). After separation, protein was transferred to Polyvinylidene difluoride membranes (Immobilon-FL) in Transfer Buffer (25mM Tris base, 192mM glycine and 20% methanol) and then, incubated with blocking buffer (Fluorescent Blocker, Thermo Fisher) for 45 min at RT. After Blocking, membranes were incubated with indicated Primary Abs (1/1000 dilution in a buffer composed by 0.1% casein, 0.2x PBS and 0.1% Tween20) for 4 hrs at room temperature (RT). Secondary Abs (1/10000 dilution in a buffer composed by 0.1% casein, 0.2x PBS, 0.1% Tween20 and 0.01% SDS) were incubated for 1 hrs at RT. Membranes were recorded with Odyssey® CLx Imaging System. Analysis and quantifications of protein expression was performed using Image Studio software (LI-COR). Data are presented as mean and error bars as standard deviation of the biological replicates. Significance was calculated using GraphPad Prism two-tailed Student's t-tests. Immunoprecipitation was performed using Thermo Fisher Scientific Protein A/G Dynabeads™, 1 µg of the specific Ab and 1000 µg of protein lysate. For endogenous co-Immunoprecipitations, beads were incubated with corresponsonding IgG as a control for specificity.

*Tandem-Ubiquitin-Binding-Entity (TUBE) assays*

Indicated cells were harvested and lysed in RIPA+ buffer (50 mM Tris-HCl pH 7.4, 1 % NP-40, 0.5 % Deoxychylate, 0.1 % SDS, 150 mM NaCl, 2 mM EDTA, 5 mM MgCl2) supplemented with Protease-Inhibitor and 1 mM DTT. To ensure protection of ubiquitin-chains, chain-unspecific non-commercial GST-Tandem-Ubiquitin-Binding-Entities (TUBEs) must be added immediately at a concentration of 100 µg/mL to lysates. 25 µg of TUBE containing lysa was boiled for 5 min at 95°C and subsequently subjected to

SDS-PAGE and Western Blotting and detection of ubiquitylated-proteins was performed with the Odyssey CLx Imaging System.

**Immunofluorescence**

Standard procedures were used for immunofluorescence (IF). For antibodies used in IF, manufacturer's manuals and instructions were used regarding concentration or buffer solutions. All primary antibodies were incubated over night at 4°C, followed by washing thrice with PBS, and subsequent incubation with the secondary antibody for 1 hour at room temperature. IF slides were imaged using FSX100 microscopy system (Olympus) and analysed using QuPath software. Or, in case of high-content microscopy, the Operetta microscope and Harmony software (Perkin Elmar) were used. For samples stained with IF, tissue-samples/ cells were counterstained with 5 µg/ml Hoechst, to highlight nuclei, for 15 minutes after secondary antibody application. Stained samples were mounted with Mowiol®40-88.

**sgRNA and shRNA Design**

sgRNAs were designed using the CRISPRtool (https://zlab.bio/guide-design-resources). shRNA sequences were designed using SPLASH-algorithm (http://splashrna.mskcc.org/)[4] or the RNAi Consortium/Broad Institute (www.broadinstitute.org/rnai-consortium/rnai-consortium-shrna-library).

**AAV and lentivirus production and purification**

Virus was packaged and synthetized in HEK293-T cells seeded in 15 cm-dishes.
For AAV production, cells (70% confluence) were transfected with the plasmid of interest (10 µg), pHelper (15 µg) and pAAV-DJ (10µg) using PEI (70 µg). After 96 hrs, the cells and medium of 3 dishes were transferred to a 50 ml Falcon tube together with 5 ml chloroform. Then, the mixture was shaken at 37°C for 60 min and NaCl (1M) was added to the mixture. After NaCl is dissolved, the tubes were centrifuged at 20,000 x g

at 4°C for 15 min and the chloroform layer was transferred to another Falcon tube together with 10 % PEG8000. As soon as the PEG800 is dissolved, the mixture was incubated at 4°C overnight and pelleted at 20,000 x g at 4°C for 15 min. The pellet was resuspended in PBS with MgCl2 and 0.001% pluronic F68, then, the virus was purified using 1X Chloroform and stored at -80C.
For Lentivirus production, HEK293-T cells (70% confluence) were transfected with the plasmid of interest (15µg), pPAX (10µg) and pPMD2 (10µg) using PEI (70µg). After 96 hrs, the medium containing lentivirus was filtered (0.45uM) and precipitated using ultracentrifugation (25,000 x g, 2.5 hrs). The virus pellet was dissolved in PBS and stored at -80°C.

**RNA-sequencing**

RNA sequencing was performed with Illumina NextSeq 500 or NextSeq 2000, as described previously [5].In brief, RNA was isolated using ReliaPrep™ RNA Cell Miniprep System Promega kit, following the manufacturer's instruction manual. mRNA was purified with NEBNext® Poly(A) mRNA Magnetic Isolation Module (NEB) and the library was generated using the NEBNext® Ultra™ II Directional RNA Library Prep Kit for Illumina, following the manufacturer's instructions. For the size-selection of the libraries, Agencourt AMPure XP Beads (Beckman Coulter) were used. Library quantification and size determination was performed using Fragment Analyzer (Agilent formerly Advanced Analytical).

**Quantification and statistical analysis**

*Analysis of publicly available data*
All publicly available data and software used for this publication are listed in the key resource table. Oncoprints were generated using cBioportal [6 7]. The following studies were used for the analysis: colorectal cancer ("COAD-TCGA, PanCancer Atlas") and GTEx colon WT samples.

Box plots using TCGA and GTEx data were generated using the online tool GEPIA [8] or GraphPad Prism software. The differential analysis was based on: "TCGA tumours vs (TCGA normal + GTEx normal)", whereas the expression data were $\log_2$(TPM+1) transformed and the $\log_2$FC was defined as median(tumour) – median(normal). p-values were calculated, if not stated otherwise, with a one-way ANOVA comparing tumour with normal tissue.

Correlation USP10 and CTNNB1 expression in tumour and normal tissue was calculated using GEPIA's software. The following datasets: "TCGA tumours" and "GTEx normal" were used for the calculation of the Spearman's correlation coefficients and significance by GEPIA's software.

KM-plot was generated using R2: Genomics Analysis and Visualization Platform, using the Tumour Colon - Smith dataset. p-values were computed using a log-rank test.

*RNA-sequencing analysis*

Fastq files were generated using Illuminas base calling software GenerateFASTQ v1.1.0.64 and overall sequencing quality was analysed using the FastQC script. Reads were aligned to the human genome (GRCh38) and murine genome (GRCm38) using Bowtie2 v2.3.2 (Langmead and Salzberg, 2012). For differential gene expression analysis, reads per gene (Ensembl gene database) were counted using the "featureCounts" package in R and non- or weakly expressed genes were removed (mean read count over all samples <6). Differentially expressed genes were called using edgeR (Robinson et al., 2010) and resulting p-values were corrected for multiple testing by false discovery rate (FDR) calculations.

*GSEA*

For gene set enrichment analyses (GSEA, (Mootha et al., 2003; Subramanian et al., 2005), normalised counts were analysed against C2, C5 and Hallmark gene sets as well as mouse-orthologue hallmark, M2 and M5 (downloaded from Molecular Signature Database hosted by Broad Institute).

Custom gene sets of cell types from the mouse small intestine were used as previously reported[9]. A cancer stem cell signature was composed of genes upregulated in LGR5+ cells versus LGR5- cells in tumours from AKP mice as reported[10].

**Mass spectrometry**

*Sample preparation*

The sample preparation was performed as described previously[11]. Briefly, lysates were precipitated by methanol/chloroform and proteins resuspended in 8 M Urea/10 mM EPPS pH 8.2. Concentration of proteins was determined by Bradford assay and 100 µg of protein per samples was used for digestion. For digestion, the samples were diluted to 1 M Urea with 10mM EPPS pH 8.2 and incubated overnight with 1:50 LysC (Wako Chemicals) and 1:100 Sequencing grade trypsin (Promega). Digests were acidified using TFA and tryptic peptides were purified by tC18 SepPak (50 mg, Waters). 125 µg peptides per sample were TMT labelled and the mixing was normalized after a single injection measurement by LC-MS/MS to equimolar ratios for each channel. 250 µg of pooled peptides were dried for offline High pH Reverse phase fractionation by HPLC.

*Offline high pH reverse phase fractionation*

Peptides were fractionated using a Dionex Ultimate 3000 analytical HPLC. 250 µg of pooled and purified TMT-labeled samples were resuspended in 10 mM ammonium-bicarbonate (ABC), 5% ACN, and separated on a 250 mm long C18 column (X-Bridge, 4.6 mm ID, 3.5 µm particle size; Waters) using a multistep gradient from 100% Solvent A (5% ACN, 10 mM ABC in water) to 60% Solvent B (90% ACN, 10 mM ABC in water) over 70 min. Eluting peptides were collected every 45 s into a total of 96 fractions, which were cross-concatenated into 12 fractions and dried for further processing.

*LC-$MS^3$ proteomics*

All mass spectrometry data was acquired in centroid mode on an Orbitrap Fusion Lumos mass spectrometer hyphenated to an easy-nLC 1200 nano HPLC system using a nanoFlex ion source (ThermoFisher Scientific) applying a spray voltage of 2.6 kV with the transfer tube heated to 300°C and a funnel RF of 30%. Internal mass calibration was enabled (lock mass 445.12003 m/z). Peptides were separated on a self-made, 32 cm long, 75µm ID fused-silica column, packed in house with 1.9 µm C18 particles (ReproSil-Pur, Dr. Maisch) and heated to 50°C using an integrated column oven (Sonation). HPLC solvents consisted of 0.1% Formic acid in water (Buffer A) and 0.1% Formic acid, 80% acetonitrile in water (Buffer B).

For total proteome analysis, a synchronous precursor selection (SPS) multi-notch MS3 method was used in order to minimize ratio compression as previously described (McAlister et al., 2014). Individual peptide fractions were eluted by a non-linear gradient from 4 to 40% B over 210 minutes followed by a step-wise increase to 95% B in 6 minutes which was held for another 9 minutes. Full scan MS spectra (350-1400 m/z) were acquired with a resolution of 120,000 at m/z 200, maximum injection time of 50 ms and AGC target value of 4 x $10^5$. The most intense precursors with a charge state between 2 and 6 per full scan were selected for fragmentation within 3 s cycle time and isolated with a quadrupole isolation window of 0.4 Th. MS2 scans were performed in the Ion trap (Turbo) using a maximum injection time of 50ms, AGC target value of 1 x $10^4$ and fragmented using CID with a normalized collision energy (NCE) of 35%. SPS-MS3 scans for quantification were performed on the 10 most intense MS2 fragment ions with an isolation window of 1.2 Th (MS) and 2 m/z (MS2). Ions were fragmented using HCD with an NCE of 65% and analyzed in the Orbitrap with a resolution of 50,000 at m/z 200, scan range of 100-200 m/z, AGC target value of 1.5 x$10^5$ and a maximum injection time of 150ms. Repeated sequencing of already acquired precursors was limited by setting a dynamic exclusion of 60 seconds and 7 ppm and advanced peak determination was deactivated.

*Quantification and Statistical Analysis*

Proteomics raw files were processed using proteome discoverer 2.2 (ThermoFisher). Spectra were recalibrated using the Homo sapiens SwissProt database (2018-11-21) and TMT as static modification at N-terminus and Lysines, together with Carbamidomethyl at cysteine residues. Spectra were searched against human database and common contaminants using Sequest HT with oxidation (M) as dynamic modification together with methionine-loss + acetylation and acetylation at the protein terminus. TMT6 (N-term, K) and carbamidomethyl were set as fixed modifications. Quantifications of spectra were rejected if average S/N values were below 5 across all channels and/or isolation interference exceeded 50%. Protein abundances were calculated by summing all peptide quantifications for each protein.

Reactome analysis were performed with PANTHER using the “Statistical overrepresentation test” tool with default settings. Proteins were considered significantly downregulated for reactome analysis when: FC<-0.5 and p-value<0.05. Heatmap visualization was performed using Morpheus (Broad Institute).

**Proximity ligation assay**

PLAs were performed according to manufacturer’s recommendation. In detail, cells were fixed with 4% PFA (EMS) diluted in PBS, and subsequently washed and permeabilized with 0.1% Triton X-100 in PBS. Blocking was performed with 5% BSA in PBS. Cells were incubated with primary antibodies against USP10 and β-Catenin in blocking solution overnight at 4 °C. The ligation reaction was carried out for 30 min, followed by washing and amplification for 2 h 30 min. For simultaneous immunofluorescence staining of the primary antibodies, the samples were incubated with secondary antibodies of respective species for 60 min in 5% BSA subsequent to completion of the PLA protocol.
Image acquisition was done using the Operetta CLS High-Content Analysis System with 40× magnification (PerkinElmer). Processing was achieved through Harmony High Content Imaging and Analysis Software (PerkinElmer) and R.

**Preparation of µSPOT Microarrays**

µSPOT peptide arrays (CelluSpots, Intavis AG, Cologne, Germany) were synthesized using a MultiPep RSi robot (Intavis AG) on in-house produced, acid labile, amino functionalized, cellulose membrane discs containing 9-fluorenylmethyloxycarbonyl-β-alanine (Fmoc-β-Ala) linkers (average loading: 131 nmol/disc – 4 mm diameter). Synthesis was initiated by Fmoc deprotection using 20% piperidine (pip) in dimethylformamide (DMF) followed by washing with DMF and ethanol (EtOH). Peptide chain elongation was achieved using a coupling solution consisting of preactivated amino acids (aas, 0.5 M) with ethyl 2-cyano-2-(hydroxyimino) acetate (oxyma, 1 M) and N,N′-diisopropylcarbodiimide (DIC, 1 M) in DMF (1:1:1, aa:oxyma:DIC). Couplings were carried out for 3×30 min, followed by capping (4% acetic anhydride in DMF) and washes with DMF and EtOH. Synthesis was finalized by deprotection with 20% pip in DMF (2×4 µL/disc for 10 min each), followed by washing with DMF and EtOH. Dried discs were transferred to 96 deep-well blocks and treated, while shaking, with sidechain deprotection solution, consisting of 90% trifluoracetic acid (TFA), 2% dichloromethane (DCM), 5% H2O and 3% triisopropylsilane (TIPS) (150 µL/well) for 1.5 h at room temperature (rt). Afterwards, the deprotection solution was removed, and the discs were solubilized overnight (ON) at rt, while shaking, using a solvation mixture containing 88.5% TFA, 4% trifluoromethanesulfonic acid (TFMSA), 5% H2O and 2.5% TIPS (250 µL/well). The resulting peptide-cellulose conjugates (PCCs) were precipitated with ice-cold ether (0.7 mL/well) and spun down at 2000×g for 10 min at 4 °C, followed by two additional washes of the formed pellet with ice-cold ether. The resulting pellets were dissolved in DMSO (250 µL/well) to give final stocks. PCC solutions were mixed 2:1 with saline-sodium citrate (SSC) buffer (150 mM NaCl, 15 mM trisodium citrate, pH 7.0) and transferred to a 384-well plate. For transfer of the PCC solutions to white coated CelluSpot blank slides (76×26 mm, Intavis AG), a SlideSpotter (Intavis AG) was used. After completion of the printing procedure, slides were left to dry ON.

*Protein expression*

β-catenin (amino acid 134-665) with an N-terminal GST-tag (GST-βcat) was expressed and purified as previously described[12].

*Microarray Binding Assay*

µSPOT slides were blocked by incubation with 2.5 mL 5% (w/v) powdered milk (Carl Roth, T245.2, MP) in PBS (pH=7.6) for 60 min at ~50 rpm and RT. Afterwards, slides were incubated with GST-βcat at 500 nM in 5% MP in 1 × PBS for 15 min before slides were washed with 3×2.5 mL 1×PBS for 1 min. To label the protein for detection, the slides were incubated with 2.5 mL of a 1:5,000 diluted, HRP-conjugated anti-GST antibody (Sigma Aldrich, RPN1236) in 5% MP in 1×PBS for 15 min, after which the slides were washed with 3×2.5 mL 1×PBS for 1 min. Peptide binding was detected through chemiluminescent detection (Lowest Sensitivity, 180s exposure time) after application of 200 µL of SuperSignal West Femto Maximum Sensitive Substrate (Thermo Scientific) per slide using a c400 imaging system (Azure).

Binding intensities were evaluated using FIJI including the Microarray Profile addon (OptiNav). After background subtraction of the mean greyscale value of the microarray surface surrounding the spots, raw greyscale intensities for each position were obtained for the left and right side of the internal duplicate on each microarray slide. Afterwards, the raw spot intensities were averaged for each individual slide, after which the mean intensities were averaged over n=3 slides used for the experiment.

**Modeling of the β-catenin/USP10 complex**

Generation of the β-catenin/USP10 complex was carried out using AlphaFold2 Multimer (AF2M) on the COSMIC2 webserver with the complete sequences of human USP10 (UniprotID Q14694.2) and human β-catenin (Uniprot ID BAG70078.1) as input. AF2M parameters were default and included the use of Amber for relaxing the predicted PDB models. Shown is the model with the highest per-residue confidence score (pLDDT).

The AF2M model of the β-catenin/USP10 complex and the β-catenin/APC complex (PDB ID: 1TH1[4] were aligned and visualized via Pymol (PyMOL Molecular Graphics System, Schrödinger, LLC.) as cartoon, surface and electrostatic representations. Side chains of interacting residues were identified using PDBePISA[13] and highlighted as stick models.

Supplementary Literature:

1. Flint Brodsly N, Bitman-Lotan E, Boico O, et al. The transcription factor Hey and nuclear lamins specify and maintain cell identity. *Elife* 2019;8 doi: 10.7554/eLife.44745 [published Online First: 2019/07/17]
2. Jiang H, Patel PH, Kohlmaier A, et al. Cytokine/Jak/Stat signaling mediates regeneration and homeostasis in the Drosophila midgut. *Cell* 2009;137(7):1343-55. doi: 10.1016/j.cell.2009.05.014 [published Online First: 2009/07/01]

3. Shaw RL, Kohlmaier A, Polesello C, et al. The Hippo pathway regulates intestinal stem cell proliferation during Drosophila adult midgut regeneration. *Development* 2010;137(24):4147-58. doi: 10.1242/dev.052506 [published Online First: 2010/11/12]
4. Pelossof R, Fairchild L, Huang CH, et al. Prediction of potent shRNAs with a sequential classification algorithm. *Nat Biotechnol* 2017;35(4):350-53. doi: 10.1038/nbt.3807 [published Online First: 2017/03/07]
5. Buchel G, Carstensen A, Mak KY, et al. Association with Aurora-A Controls N-MYC-Dependent Promoter Escape and Pause Release of RNA Polymerase II during the Cell Cycle. *Cell Rep* 2017;21(12):3483-97. doi: 10.1016/j.celrep.2017.11.090 [published Online First: 2017/12/21]
6. Gao J, Aksoy BA, Dogrusoz U, et al. Integrative analysis of complex cancer genomics and clinical profiles using the cBioPortal. *Sci Signal* 2013;6(269):pl1. doi: 10.1126/scisignal.2004088 [published Online First: 2013/04/04]
7. Cerami E, Gao J, Dogrusoz U, et al. The cBio cancer genomics portal: an open platform for exploring multidimensional cancer genomics data. *Cancer Discov* 2012;2(5):401-4. doi: 10.1158/2159-8290.CD-12-0095 [published Online First: 2012/05/17]
8. Tang Z, Li C, Kang B, et al. GEPIA: a web server for cancer and normal gene expression profiling and interactive analyses. *Nucleic Acids Res* 2017;45(W1):W98-W102. doi: 10.1093/nar/gkx247 [published Online First: 2017/04/14]
9. Haber AL, Biton M, Rogel N, et al. A single-cell survey of the small intestinal epithelium. *Nature* 2017;551(7680):333-39. doi: 10.1038/nature24489 [published Online First: 2017/11/17]
10. Fumagalli A, Oost KC, Kester L, et al. Plasticity of Lgr5-Negative Cancer Cells Drives Metastasis in Colorectal Cancer. *Cell Stem Cell* 2020;26(4):569-78 e7. doi: 10.1016/j.stem.2020.02.008 [published Online First: 2020/03/15]
11. Klann K, Tascher G, Munch C. Functional Translatome Proteomics Reveal Converging and Dose-Dependent Regulation by mTORC1 and eIF2alpha. *Mol Cell* 2020;77(4):913-25 e4. doi: 10.1016/j.molcel.2019.11.010 [published Online First: 2019/12/10]
12. Wendt M, Bellavita R, Gerber A, et al. Bicyclic beta-Sheet Mimetics that Target the Transcriptional Coactivator beta-Catenin and Inhibit Wnt Signaling. *Angew Chem Int Ed Engl* 2021;60(25):13937-44. doi: 10.1002/anie.202102082 [published Online First: 2021/03/31]
13. Krissinel E, Henrick K. Inference of macromolecular assemblies from crystalline state. *J Mol Biol* 2007;372(3):774-97. doi: 10.1016/j.jmb.2007.05.022 [published Online First: 2007/08/08]
